# Human-chimpanzee tetraploid system defines mechanisms of species-specific neural gene regulation

**DOI:** 10.1101/2025.03.31.646367

**Authors:** Janet H.T. Song, Ava C. Carter, Evan M. Bushinsky, Samantha G. Beck, Jillian E. Petrocelli, Gabriel T. Koreman, Juliana Babu, David M. Kingsley, Michael E. Greenberg, Christopher A. Walsh

## Abstract

A major challenge in human evolutionary biology is to pinpoint genetic differences that underlie human-specific traits, such as increased neuron number and differences in cognitive behaviors. We used human-chimpanzee tetraploid cells to distinguish gene expression changes due to *cis*-acting sequence variants that change local gene regulation, from *trans* expression changes due to species differences in the cellular environment. In neural progenitor cells, examination of both *cis* and *trans* changes — combined with CRISPR inhibition and transcription factor motif analyses — identified *cis*-acting, species-specific gene regulatory changes, including to *TNIK, FOSL2*, and *MAZ*, with widespread *trans* effects on neurogenesis-related gene programs. In excitatory neurons, we identified *POU3F2* as a key *cis*-regulated gene with *trans* effects on synaptic gene expression and neuronal firing. This study identifies *cis*-acting genomic changes that cause cascading *trans* gene regulatory effects to contribute to human neural specializations, and provides a general framework for discovering genetic differences underlying human traits.

## Introduction

Compared to chimpanzees, humans have a three-fold larger brain, a prolonged period of neuronal maturation, and differences in cognitive and social behaviors (Pollen et al., 2023; Vanderhaeghen and Polleux, 2023). However, the genetic changes between humans and chimpanzees that underlie these phenotypic differences remain poorly understood. Prior studies in vertebrates suggest that the vast majority of evolutionary differences between species are due to context-specific gene expression changes that are mediated by non-coding sequence variation at gene regulatory elements (King and Wilson, 1975; Carroll, 2008; McLean et al., 2011; Jones et al., 2012), but identifying these key genomic changes is challenging due to the immense spatiotemporal complexity of the gene regulatory landscape.

A common approach to begin to identify gene regulatory changes underpinning human evolution is to directly compare cultured or primary neural cell types between human and non-human primates. This approach has identified hundreds to thousands of genes that are differentially expressed in specific cell types, brain regions, and developmental time points (Ma et al., 2022; Caglayan et al., 2023; Jorstad et al., 2023). Although these studies provide an upper bound on the gene expression changes that may be causal for human-specific phenotypes in particular regulatory contexts, identifying the key molecular changes from among these candidate genes and linking them to their causative regulatory sequence variants remains a daunting challenge. This is further complicated by the paucity of available primate fetal samples and the challenge of accurately matching developmental stage and cell type identity across species.

The recent generation of human-chimpanzee tetraploid induced pluripotent stem cells (iPSCs) provides an orthogonal approach to narrow this search space (Agoglia et al., 2021; Song et al., 2021; Pavlovic et al., 2022). By examining tetraploid cells that contain both the human and chimpanzee genome and comparing them to cells that contain each genome separately (what we collectively term the “human-chimpanzee tetraploid system”), we can distinguish species-differential genes that are regulated in *cis* — due to a nearby linked sequence variant — from those that are regulated in *trans* simply due to changes in the cellular environment, such as the levels of transcription factors (TFs) (Fig. 1A). Although this system has immense promise to identify key genetic changes governing human evolution, the human-chimpanzee tetraploid system has only been used to identify and experimentally model *cis*-regulated genes to date (Agoglia et al., 2021; Gokhman et al., 2021), leaving it an open question whether this system can actually connect *cis*-regulated gene expression changes to their causative sequence variants in practice. In addition, because *cis* changes with large effect sizes likely have cascading *trans* effects on the expression of many genes, we hypothesized that simultaneously examining <u>both</u> *cis*- and *trans*-regulated genes may substantially boost our power to identify *cis* changes with amplified *trans*-acting gene regulatory effects as key contributors to human evolution.

**Figure 1:**
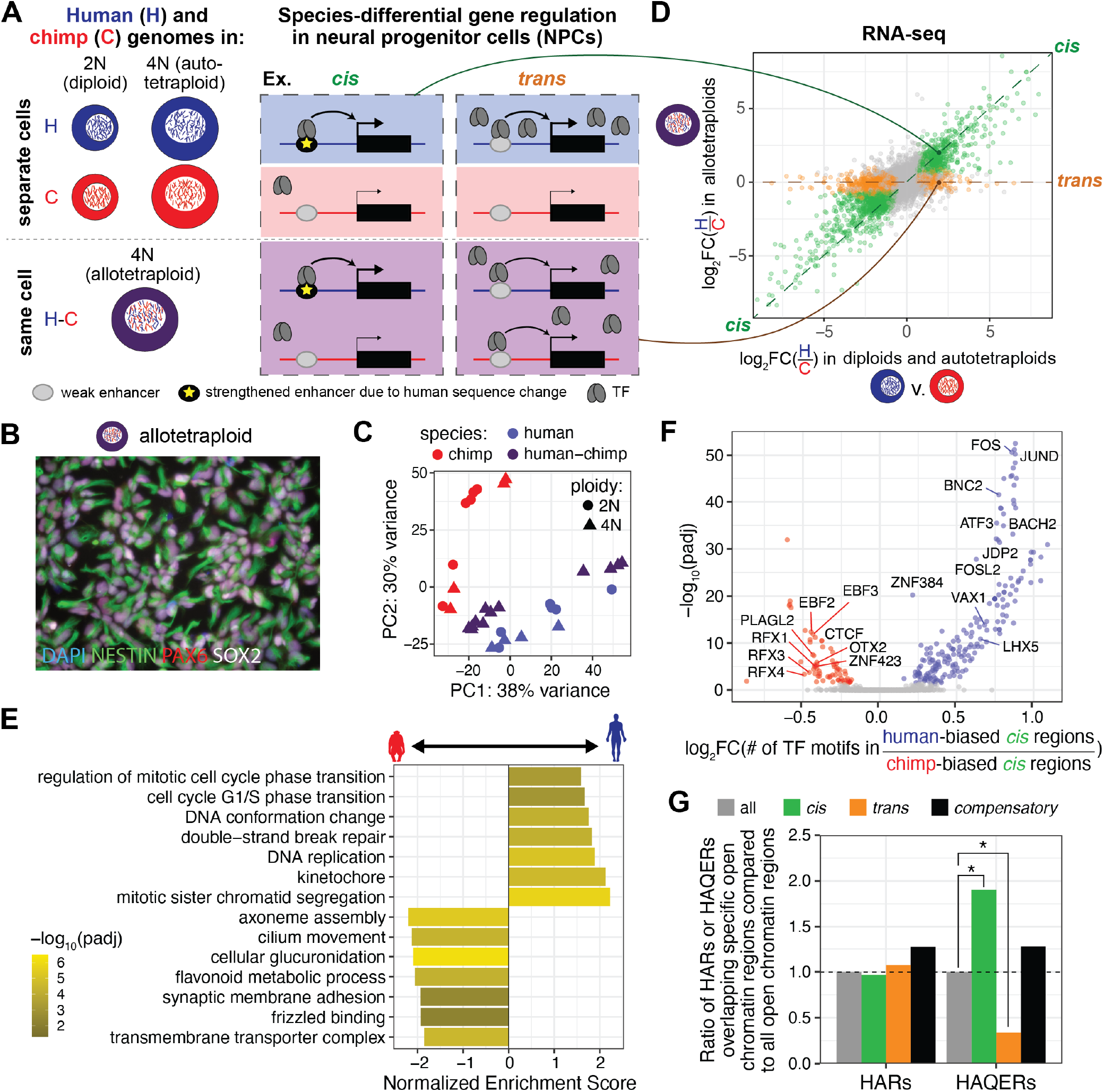
Distinguishing *cis* and *trans* gene expression and chromatin accessibility changes between human and chimpanzee NPCs. (A) Schematic of potential mechanisms for *cis* and *trans* changes. (B) Representative image of allotetraploid NPCs stained for PAX6, NESTIN, and SOX2. (C) PCA of RNA-seq shows separation by species. (D) RNA-seq identifies *cis*-regulated (green) and *trans*-regulated (orange) gene expression changes. (E) Representative, significant terms from gene-set enrichment analysis of differential genes between human and chimpanzee NPCs. (F) TF motifs enriched in *cis*-regulated open chromatin regions. (G) Ratio of the proportion of HARs or HAQERs overlapping all, *cis*-regulated, *trans*-regulated, or *compensatory* open chromatin regions compared to the proportion of HARs or HAQERs overlapping all open chromatin regions. *: *pad j* < 0.05; **: *pad j* < 0.01; ****: *pad j* < 0.0001. See also Fig. S1, Tables S1 and S2, and Materials and Methods.

In this study, we applied the human-chimpanzee tetraploid system to iPSC-derived neural progenitor cells (NPCs), which model the NPCs that have expanded in proliferative capacity to result in the increased neuron number and brain size characteristic of humans (Pollen et al., 2023). Increased neuron number in humans is likely due, at least in part, to increased self-renewal and delayed terminal differentiation of NPCs upon mitosis (Dehay and Huttner, 2024). Although comparative genomics and transcriptomics efforts to date have identified a handful of evolutionary differences that modulate this process (Florio et al., 2015; Fiddes et al., 2018; Suzuki et al., 2018; Liu et al., 2024), the genome-wide regulatory changes that govern the evolutionary expansion of neuron number in humans are still largely uncharacterized.

Using the human-chimpanzee tetraploid system, we distinguished *cis*- and *trans*-regulated genes and open chromatin regions in iPSC-derived NPCs. We used a CRISPR inhibition (CRISPRi) screen to test whether *cis*-regulated open chromatin regions regulate nearby *cis*-regulated target genes, and performed TF motif analyses of *trans*-regulated open chromatin regions to identify *cis*-regulated TFs that have widespread *trans* effects. These include a *cis*-acting regulatory region near *TNIK* and two *cis*-regulated TFs, FOSL2 and MAZ, with substantial *trans* effects on neurogenesis-related gene programs. We also extended these approaches to excitatory neurons, a major building block of neuronal circuits in the brain, and identified POU3F2 as a key *cis*-regulated TF with strong *trans* effects on synaptic gene expression and neuronal firing. Our results demonstrate how using the human-chimpanzee tetraploid system to examine both *cis* and *trans* effects can lead to a genomewide understanding of human-specific gene regulatory changes.

## Results

### Distinguishing cis- and trans-regulated genes in neural progenitor cells (NPCs)

To identify genetic changes that underlie human-chimpanzee differences in NPCs, we differentiated three human diploid, three chimpanzee diploid, two human autotetraploid, two chimpanzee autotetraploid, and six human-chimpanzee allotetraploid iPSC lines for 14 days into NPCs (Fig. 1A, B; Fig. S1A; Materials and Methods). Autotetraploid lines are derived from the fusion of two diploid iPSCs of the same species and serve as tetraploidization controls, while allotetraploid cells are derived from the fusion of a human iPSC line and a chimpanzee iPSC line (Song et al., 2021). We performed RNA-seq to identify species-differential genes and ATAC-seq to identify species-differential chromatin accessibility for two biological replicates per line. Open chromatin regions identified by ATAC-seq are associated with regulatory activity (Gasperini et al., 2020).

As expected, principal component analysis (PCA) separated human from chimpanzee samples along the first two principal components (PCs) of variation for RNA-seq and along the first PC for ATAC-seq, with human-chimpanzee samples falling in between the human and chimpanzee samples respectively (Fig. 1C; Fig. S1B). All lines generated homogenous populations of PAX6+, NESTIN+, SOX2+ NPCs by immunofluorescence and expressed dorsal forebrain NPC markers by RNA-seq (Fig. 1B; Fig. S1A, C). There were some differences in NPC morphology and the relative strength of marker gene expression between lines (Fig. S1A, C), but these differences were not correlated with species or ploidy, suggesting that they represent normal variability in differentiations between different iPSC lines (Kyttälä et al., 2016). Diploid and autotetraploid NPCs were well mixed by PCA within each species (Fig. 1C; Fig. S1B), and only 11 genes were differentially expressed between diploids and autotetraploids in both human and chimpanzee NPCs (Table S1), suggesting that tetraploidization does not lead to overt gene expression changes in NPCs.

To identify and distinguish *cis*- from *trans*-regulated genes, we performed differential gene expression analysis between human and chimpanzee NPCs and allele-specific expression analysis between human and chimpanzee alleles within allotetraploids. *Cis*-regulated genes differ in expression due to nearby sequence variants on the same DNA molecule (Fig. 1A). Because *cis*-regulated gene expression changes are due to sequence variants and not due to changes in the cellular environment, they have both significant differential expression when the human and chimpanzee orthologs are in separate cellular environments (diploids and autotetraploids) and significant allele-specific expression when the human and chimpanzee orthologs are in the same cellular environment (allotetraploids). *Trans*-regulated genes differ in expression due to changes in the cellular environment, such as different levels of TFs, chromatin remodelers, and other molecules, and not due to linked sequence changes. Accordingly, they have significant differential expression between human and chimpanzee orthologs in separate cellular environments (diploids and autotetraploids) but nonsignificant allele-specific expression between human and chimpanzee orthologs exposed to the same cellular environment (allotetraploids). We note that although having both significant differential expression and significant allele-specific expression indicates that a gene is regulated at least in part due to a *cis* contribution, these gene expression differences may also have a *trans* contribution. For simplicity, we refer to genes with any *cis* contribution as *cis*-regulated genes, and genes with only a *trans* contribution as *trans*-regulated genes.

There are 2,181 *cis*-regulated genes and 1,574 *trans*-regulated genes between human and chimpanzee NPCs at a fold-change > 1.5 (Fig. 1D; Table S1). *Cis*- and *trans*-regulated genes with increased expression in human NPCs compared to chimpanzee NPCs (human-biased) are enriched for gene ontology terms related to the cell cycle, consistent with prolonged neurogenesis in the human lineage (Fig. 1E; Fig. S1D; Table S1). These include genes such as *ANKRD53, TTN*, and *NEK6* that are required for mitosis (Liu and Meinke, 1998; Yin et al., 2003; Kim and Jang, 2016). In contrast, *cis*- and *trans*-regulated genes with decreased expression in human NPCs compared to chimpanzee NPCs (chimpanzee-biased) are enriched for synaptic and metabolic terms consistent with neuronal differentiation. Notably, the anticipatory expression of neuronal gene programs, including synaptic genes, has been observed in NPCs that preferentially give rise to terminally differentiated neurons at the next cell division (Zahr et al., 2018; Polioudakis et al., 2019), and changes in metabolism have been linked to NPC fate decisions (Iwata and Vanderhaeghen, 2021). This suggests that chimpanzee NPCs may be more transcriptionally primed for terminal differentiation compared to human NPCs. Taken together, these enrichments parallel the expected molecular difference between human and chimpanzee NPCs *in vivo*, where human NPCs undergo additional cell divisions prior to terminal differentiation into neurons in order to produce a larger number of neurons. This suggests that the human-chimpanzee tetraploid system can be used to uncover physiologically-relevant biology in NPCs that drive these species differences.

### Identifying cis- and trans-regulated open chromatin regions in NPCs

To uncover regulatory elements that might drive species-differential gene expression, we performed ATAC-seq and identified 12,372 *cis*-regulated open chromatin regions and 18,509 *trans*-regulated open chromatin regions between human and chimpanzee NPCs (Fig. S1E; Table S2). Open chromatin regions with differences in accessibility in human compared to chimpanzee NPCs are enriched near genes involved in mitochondria and metabolism, consistent with the key role mitochondria play in NPC fate decisions (Iwata and Vanderhaeghen, 2021), as well as near genes related to splicing factors and forebrain formation (Fig. S1F; Table S2).

Transcription factor (TF) motif analyses of *cis*-regulated open chromatin regions identified AP-1 motifs at human-biased regions and motifs for EBF and RFX TFs at chimpanzee-biased regions (Fig. 1F; Table S2). Although little is known about the role of TFs that bind the AP-1 motif in NPCs, both EBF and RFX TFs have been shown to promote the terminal differentiation of NPCs into neurons (Garcia-Dominguez et al., 2003; Choi et al., 2024). Decreased binding of EBF and RFX TFs in human NPCs would be consistent with epigenomic rewiring between humans and chimpanzees to shift the balance between NPC self-renewal and terminal differentiation toward self-renewal in human NPCs.

### HAQERs but not HARs are enriched at cis-regulated open chromatin regions

Distinguishing *cis* and *trans* changes allows us to infer the functional effects of previously identified genomic regions with signatures of positive selection in the human lineage. These include (1) human accelerated regions (HARs), characterized by strong conservation through non-human mammals coupled with accelerated substitution rates specifically in humans (Girskis et al., 2021; Keough et al., 2023; Bi et al., 2023); and (2) human ancestor quickly evolved regions (HAQERs), which have the highest substitution rates in the human genome and likely represent novel human regulatory elements (Mangan et al., 2022). Although HARs and HAQERs are enriched near genes expressed in the brain, relatively few of these elements have been shown to regulate gene expression endogenously (Aldea et al., 2021; Dutrow et al., 2022; Liu et al., 2024), raising the question of whether these elements are enriched for functional regulatory effects in their genomic context.

If HARs or HAQERs have regulatory effects due to internal sequence changes, they will be enriched among *cis*-regulated open chromatin regions and depleted from *trans*-regulated open chromatin regions. However, if sequence variants in HARs or HAQERs merely act to maintain the same level of chromatin accessibility in reaction to changes in the *trans* environment, then we might observe an enrichment at *compensatory* open chromatin regions. *Compensatory* open chromatin regions have significant allele-specific accessibility in the same cellular environment (allotetraploids) but are not differentially accessible in separate cellular environments (diploids and autotetraploids), indicating that they are affected by *cis* and *trans* effects that act in opposing directions (Fig. S1E).

Surprisingly, HARs are not enriched for *cis*-regulated open chromatin regions (Fig. 1G), suggesting that internal sequence variants in HARs do not generally produce changes in regulatory activity. To determine whether this lack of enrichment for HARs at *cis*-regulated open chromatin regions is due to differences in genome quality and availability across studies, we separately examined all HAR sets identified prior to 2021 (Girskis et al., 2021), zooHARs identified using 241 mammalian genomes (Keough et al., 2023), and lineage HARs identified using a densely sampled primate tree (Bi et al., 2023). None of these HAR subsets were significantly enriched for any gene regulatory subset of open chromatin regions, although the proportion of *compensatory* open chromatin regions was consistently higher than *cis*-regulated open chromatin regions across all subsets (Fig. S1G). In contrast, HAQERs did show significant enrichment among *cis*-regulated open chromatin regions (adjusted *p* = 0.03) and significant depletion from *trans*-regulated open chromatin regions (adjusted *p* = 0.03), without a significant difference in *compensatory* open chromatin regions (adjusted *p* = 0.65) (Fig. 1G). This suggests that novel regulatory elements like HAQERs include a larger proportion of sites that have species-differential, *cis* effects compared to sequence changes in highly conserved elements like HARs.

### CRISPR inhibition screen identifies seven cis-regulated enhancers

Although DNA sequence comparisons have identified candidate species-specific regulatory elements like HARs and HAQERs, only a tiny fraction of these elements have been shown to regulate gene expression endogenously (Baumgartner et al., 2024) and many may lack sequence-intrinsic activity (Fig. 1G), motivating us to determine whether the human-chimpanzee tetraploid system can provide an orthogonal, complementary approach to nominate and identify causative regulatory changes. We examined *cis*-regulated open chromatin regions with a CRISPR inhibition (CRISPRi) screen, which uses a catalytically inert Cas9 that is fused to the KRAB repressor domain (dCas9-KRAB) to heterochromatinize target regions (Gilbert et al., 2014). Because prior studies suggest that most human-chimpanzee sequence variants — including ones with genomic signatures of selection — are either non-functional or have extremely small effect sizes (Fair and Pollen, 2023), a CRISPRi screen is a potentially high-throughput approach to quickly identify *cis*-regulated open chromatin regions with large effects on gene expression for additional downstream studies.

Of the >12,000 *cis*-regulated open chromatin regions, we tested those that met several technical and biological criteria in our CRISPRi screen (Fig. 2A; Materials and Methods). First, we chose *cis*-regulated open chromatin regions >2 kb from the nearest transcription start site and >1 kb from the nearest exon to avoid heterochromatinization of coding regions. Second, we required that nearby *cis*-regulated target genes would be expressed sufficiently highly to detect a >25% change in expression by single cell RNA-seq (scRNA-seq). To capture distal *cis* regulation, we used a broad definition to associate open chromatin regions with nearby genes. Nearby *cis*-regulated genes were defined as those within the same topologically-associated domain (TAD) (Won et al., 2016; Schmitt et al., 2016; Bonev et al., 2017) or within 500 kb of the *cis*-regulated open chromatin region.

**Figure 2:**
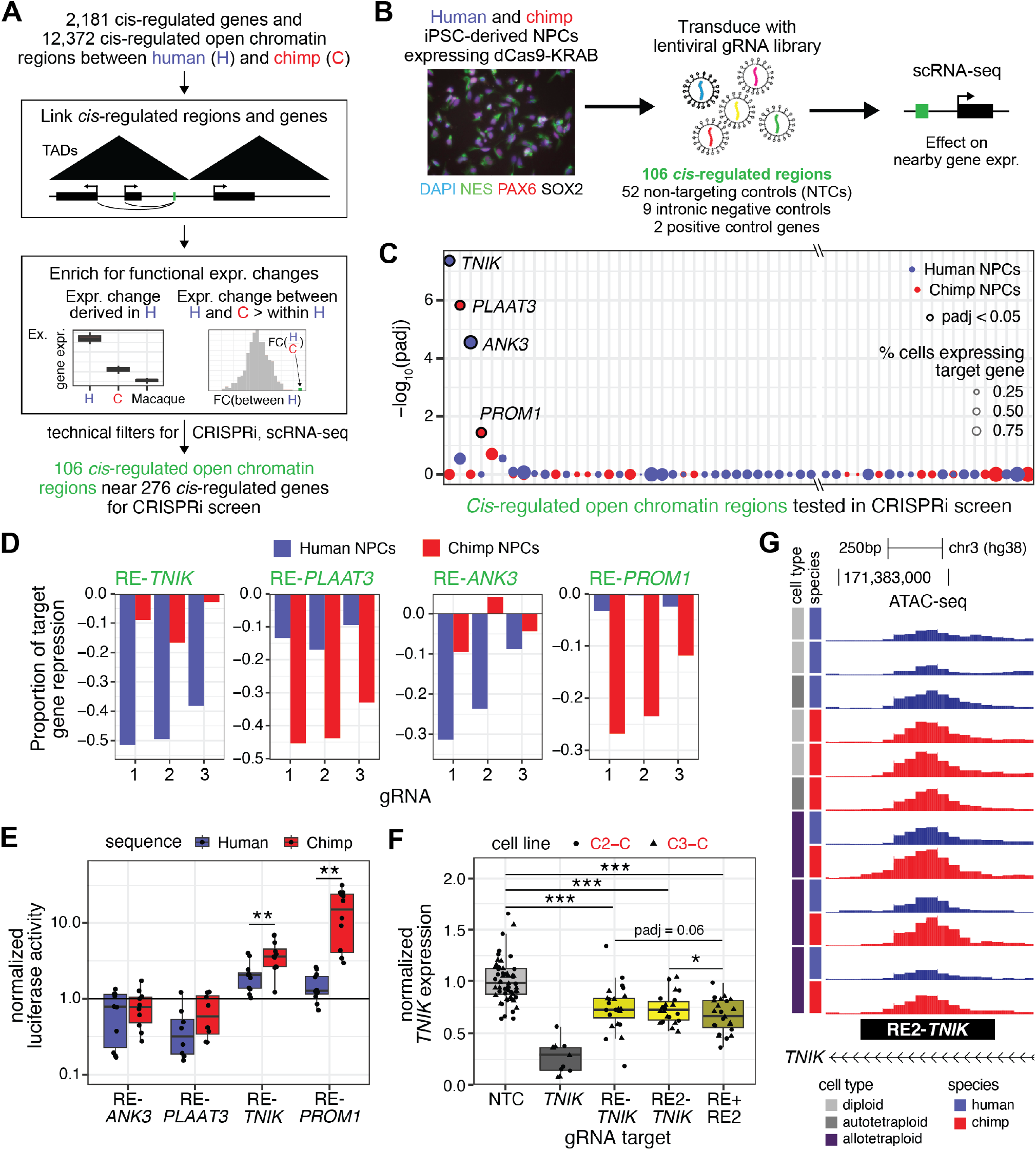
CRISPR inhibition screen in NPCs identifies *cis*-regulated enhancers of nearby genes. (A) Workflow to identify *cis*-regulated open chromatin regions for CRISPRi screen. (B) Schematic of CRISPRi screen. Image shows NESTIN+, PAX6+, SOX2+ NPCs differentiated from the chimpanzee C3-C line. (C) Adjusted p-values for the effect of CRISPRi targeting each *cis*-regulated open chromatin region on a nearby *cis*-regulated gene in human NPCs (blue) or chimpanzee NPCs (red). Significant hits are outlined in black and labeled with their target gene. Only a subset of nonsignificant tested regions are shown. (D) Proportion of target gene repression in the CRISPRi screen for each gRNA targeting RE-*TNIK*, RE-*PLAAT3*, RE-*ANK3*, and RE-*PROM1* in human and chimpanzee NPCs. (E) Luciferase enhancer activity of the human and chimpanzee sequences of RE-*TNIK*, RE-*PLAAT3*, RE-*ANK3*, and RE-*PROM1*. (F) qPCR for *TNIK* expression in NPCs differentiated from two chimpanzee lines (C2-C and C3-C) and infected with a combination of a NTC gRNA or gRNAs targeting the *TNIK* promoter, RE-*TNIK*, or RE2-*TNIK* (Materials and Methods). Note that statistical analysis was performed with data from both chimpanzee lines and human lines (Fig. S2E). (G) Read pile-ups from ATAC-seq for representative diploid, autotetraploid, and allotetraploid NPCs at RE2-*TNIK*. *: *pad j* < 0.05; **: *pad j* < 0.01; ***: *pad j* < 0.001. See also Fig. S2, Fig. S3, Table S3, and Materials and Methods.

To enrich for regulatory changes that are derived in humans and likely to have downstream functional effects, we further prioritized *cis*-regulated open chromatin regions for those that met several biological criteria (detailed in Materials and Methods), including the following: To examine human-derived changes, we filtered for *cis*-regulated open chromatin regions that contain 1+ base pairs that are conserved through macaques, gorillas, and chimpanzees but change in humans, and that are near 1+ *cis*-regulated genes where the direction of the gene expression change is consistent between humans and chimpanzees and humans and macaques (Zhu et al., 2018). We also enriched for changes that are likely to have downstream *trans* effects by selecting *cis*-regulated open chromatin regions near 1+ *cis*-regulated genes that are implicated in neurodevelopmental disorders (Firth et al., 2009; Leblond et al., 2021) or that are intolerant to heterozygous loss-of-function mutations in humans (Lek et al., 2016), suggesting that modest changes in expression may have neurological consequences. In addition, we required that *cis*-regulated open chromatin regions are near 1+ *cis*-regulated genes where the expression change between humans and chimpanzees is significantly larger than the variation in expression between modern, healthy humans (Starr et al., 2023), indicating that the human-chimpanzee difference in expression may be functional.

The resulting set of 106 *cis*-regulated open chromatin regions were each targeted by three guide RNAs (gRNAs) matching sequences identical in humans and chimpanzees (Table S3). Each region is near 1™ 7 potential *cis*-regulated target genes (median = 1 gene, mean = 2 genes) (Fig. S2A), underscoring the need to functionally test these regions within their genomic context to accurately identify target genes. One human and one chimpanzee iPSC line that endogenously express dCas9-KRAB were differentiated into NPCs and infected with a lentivirus pool containing non-targeting control (NTC) gRNAs, negative control gRNAs targeting intronic regions, positive control gR-NAs targeting gene promoters, and gRNAs targeting the 106 *cis*-regulated open chromatin regions. scRNA-seq was performed seven days after infection (Fig. 2B). As expected, positive control gRNAs targeting gene promoters reduced target gene expression, and negative control gRNAs targeting intronic regions within 1™ 5 kb of the exons of three different genes had no effect on nearby gene expression (Fig. S2B, C; Table S3).

CRISPRi targeting of 7 of the 106 *cis*-regulated open chromatin regions decreased expression of a nearby gene (Fig. 2C; Fig. S2B; Table S3). Three of these regions, including one region that overlapped a HAR, affected genes that were not *cis*-regulated (Fig. S2B), suggesting that their *cis* effects on gene expression are compensated by other *cis* or *trans* factors. Of the four regions that affected *cis*-regulated genes (Fig. 2C), two are human-biased and regulated nearby human-biased, *cis*-regulated genes: *ANK3*, which encodes a scaffolding protein associated with bipolar disorder (Stahl et al., 2019), and *PLAAT3*, which encodes a phospholipid-modifying enzyme, respectively. The other two regions are chimpanzee-biased and regulated nearby chimpanzee-biased genes: *TNIK*, which encodes a serine/threonine kinase, and *PROM1*, which encodes a transmembrane protein implicated in hippocampal neurogenesis (Kempermann et al., 2006), respectively. For clarity, we labeled these open chromatin regions as regulatory elements (REs) of their target genes (e.g. RE-*ANK3*). *ANK3, TNIK*, and *PROM1* modulate Wnt signaling (Shitashige et al., 2010; Coba et al., 2012; Durak et al., 2015; Lee et al., 2017; Bahn and Ko, 2023), a major pathway in neurogenesis (Chenn and Walsh, 2002), and mutations in *ANK3, TNIK*, and *PLAAT3* have all been linked to neurological deficits (Durak et al., 2015; Anazi et al., 2016; Schuermans et al., 2023). This suggests that changes in the regulatory activities of these *cis*-regulated open chromatin regions may have contributed to changes in neurodevelopment between human and chimpanzee NPCs. More broadly, our hit rate in the CRISPRi screen is comparable to prior screens of regions selected by genomic sequence comparison (Fair et al., 2023; Geller et al., 2024), while also simultaneously filtering for hits that specifically affect *cis*-regulated genes. These results demonstrate the utility of the human-chimpanzee tetraploid system to prospectively identify regulatory changes with *cis* effects on nearby species-specific gene expression.

### Multiple enhancers contribute to species-differential TNIK expression

Of the four *cis*-regulated open chromatin regions that affected *cis*-regulated genes in the CRISPRi screen, CRISPRi of RE*ANK3* and RE-*PROM1* significantly decreased expression of their target genes only in the species where these REs are more accessible and the target genes are more highly expressed, as expected. In contrast, CRISPRi of RE-*TNIK* and RE-*PLAAT3* significantly decreased expression only in the species where these REs are less accessible and their target genes are less expressed (Fig. 2C, D). The discrepancy between species bias in chromatin accessibility and CRISPRi targeting results for RE*TNIK* and RE-*PLAAT3* may indicate that differences in chromatin accessibility are not reflective of enhancer strength or that additional regulatory elements near their target genes buffer the endogenous effects of these REs in the species with higher target gene expression. We further tested these possibilities.

Luciferase enhancer reporter assays revealed that species bias in chromatin accessibility accurately reflects species bias in enhancer strength. The chimpanzee sequences of RE-*TNIK* and RE-*PROM1* drove significantly higher luciferase activity than their human counterparts, consistent with their species bias in chromatin accessibility (Fig. 2E). These differences in enhancer activity were observed when sequences were tested in either human or chimpanzee NPCs, confirming that these differences are *cis*-regulated (Fig. S2D). For RE-*ANK3* and RE-*PLAAT3*, we could not assess relative enhancer strength because these REs did not show enhancer activity in our assays (Fig. 2E). It is likely that their regulatory activity requires additional sequence elements not captured in our reporter constructs.

To test whether other regulatory elements near RE-*TNIK* buffer its effect, we examined RE2-*TNIK*, an open chromatin region at the TNIK locus that scored just above the significance threshold for *cis*-regulation (Fig. 2F, G; Fig. 3A; Table S2). RE-*TNIK* and RE-*TNIK2* are ∼4 kb apart and are both located in the first intron of *TNIK*, ∼80 kb from the *TNIK* promoter and ∼10 kb from the second exon of *TNIK*. We find that the chimpanzee RE2-*TNIK* sequence had stronger enhancer activity than the human RE2-*TNIK* sequence in luciferase assays when tested in either human or chimpanzee NPCs, establishing it as a *cis*-regulated enhancer (Fig. S2D). Similar to RE-*TNIK*, CRISPRi of RE2-*TNIK* in human NPCs had a stronger effect on *TNIK* expression than CRISPRi in chimpanzee NPCs (Fig. 2F; Fig. S2E). Targeting RE-*TNIK* and RE2-*TNIK* together within the same cells decreased *TNIK* expression more than targeting either RE-*TNIK* alone (adjusted *p* = 0.064) or RE2-*TNIK* alone (adjusted *p* = 0.0135), although the change in effect size was far from additive (Fig. 2F; Fig. S2E). These results indicate that *TNIK* is regulated by at least two species-differential enhancers and that there are likely additional elements or mechanisms that buffer expression at the *TNIK* locus. Taken together, weaker knockdown of *TNIK*, a chimpanzee-biased gene, when targeting RE-*TNIK* in chimpanzee NPCs is not due to discordance between enhancer strength and chromatin accessibility, but likely reflects buffering by additional mechanisms in chimpanzee NPCs at this locus. Gene expression buffering by multiple regulatory elements may also explain why only a minority of highly-filtered open chromatin regions affect nearby gene expression in our CRISPRi screen and others (Gasperini et al., 2019).

**Figure 3:**
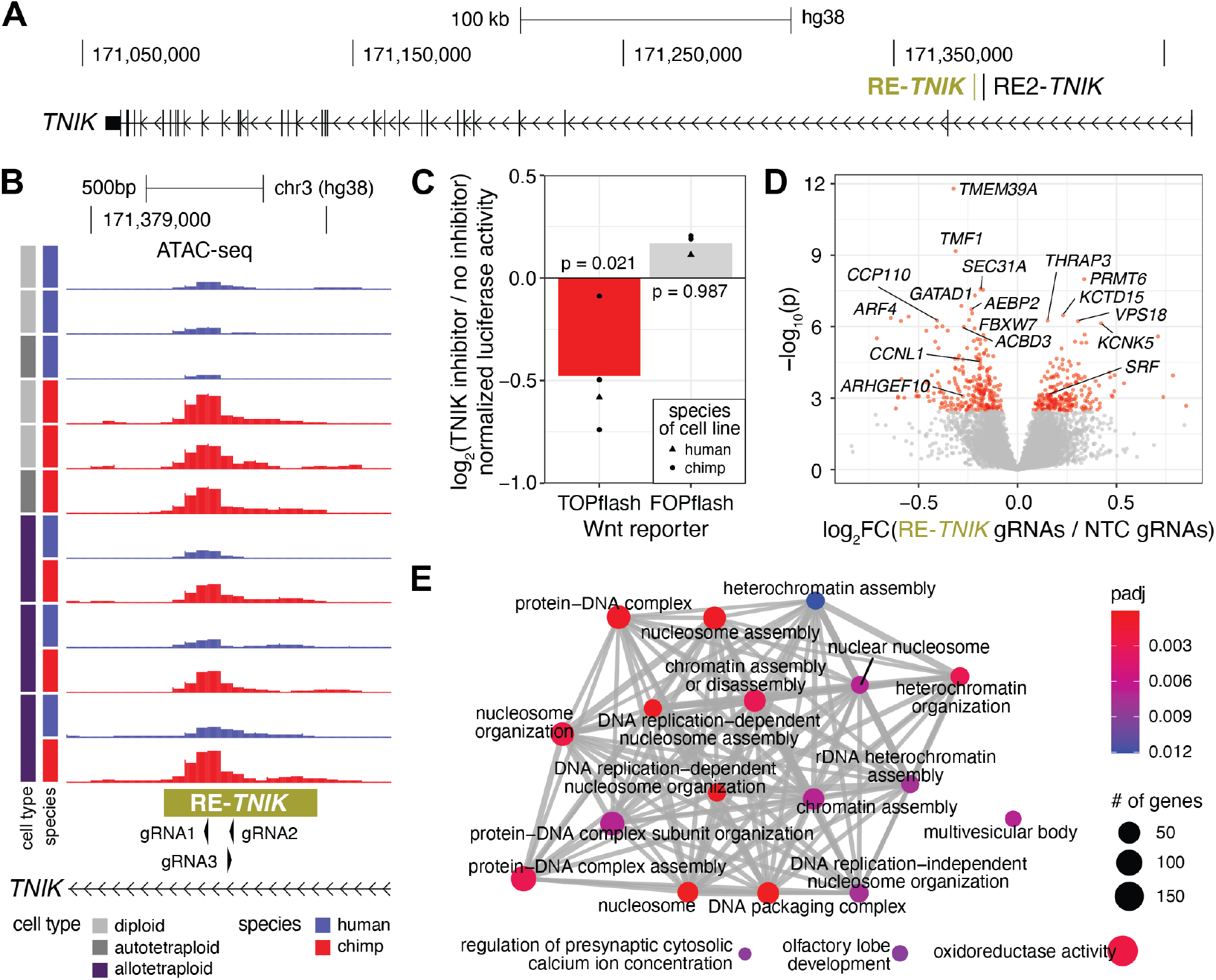
RE-*TNIK* affects cell cycle gene expression in NPCs. (A) Genomic interval containing *TNIK* and RE-*TNIK* in hg38 coordinates. (B) RE-*TNIK* is a *cis*-regulated open chromatin region with increased chromatin accessibility in chimpanzee compared to human NPCs. The location of the three gRNAs used to target RE-*TNIK* are indicated. (C) 50µM of KY-05009, a TNIK inhibitor, decreased luciferase activity from the Wnt reporter TOPflash in the presence of 50µM of CHIR-99021, a Wnt agonist (red). KY-05009 had no effect on luciferase activity from the control reporter FOPflash (gray). (D) Volcano plot of differential gene expression between chimpanzee NPCs infected with gRNAs targeting RE-*TNIK* and NTC gRNAs. Genes significant at 5% FDR are in red. Highly significant genes, as well as examples of genes associated with Wnt signaling, are labelled. (E) Significant terms from gene-set enrichment analysis of differential gene expression between chimpanzee NPCs infected with gRNAs targeting RE-*TNIK* and NTC gRNAs. See also Fig. S4, Table S4, and Materials and Methods.

Regulatory element redundancy is likely a widespread mechanism to buffer gene expression changes across species. Although *cis*-regulated open chromatin regions that are human-biased but not chimpanzee-biased are enriched near human-biased *cis*-regulated genes and vice versa, the magnitude of these enrichments are modest (Fig. S3A), suggesting that multiple regulatory changes may be required for a gene expression change. Indeed, there is a significantly higher proportion of *cis*-regulated open chromatin regions that are human-biased but not chimpanzee-biased near human-biased, *cis*-regulated genes (Fig. S3B). For example, *PLAAT3* is flanked by four human-biased, *cis*-regulated open chromatin regions including RE-*PLAAT3*, suggesting that evolutionary sequence variants in multiple regulatory elements may contribute to its species-differential expression (Fig. S3C). Similar enrichments are also observed for other gene regulation categories and directions of effect (Fig. S3B). Because genes with decreased expression variation due to genomic redundancies are enriched for those that perform essential functions (Zhou et al., 2008; Wolf et al., 2023), changes in *TNIK* and *PLAAT3* expression between human and chimpanzee NPCs may have resulted in significant functional effects.

### RE-TNIK may affect Wnt signaling and cell cycle gene expression in NPCs

Several lines of evidence suggest that changes to *TNIK* expression levels might contribute to biological differences between human and chimpanzee NPCs. *TNIK* is *cis*-regulated, with higher gene expression and protein levels in chimpanzee compared to human NPCs (Fig. S4A, B). *TNIK* is highly constrained (pLI = 1) (Lek et al., 2016), suggesting that changes to *TNIK* expression are likely to have phenotypic effects. It encodes a serine/threonine kinase with Wnt-dependent and Wnt-independent functions in neurons and in cancer cell lines (Shitashige et al., 2010; Coba et al., 2012; Lee et al., 2017). *TNIK* null mutations in humans are associated with intellectual disability without gross defects in brain size (Anazi et al., 2016), and mouse *TNIK* knock-outs have many neurological deficits, including impaired cognitive behaviors (Coba et al., 2012; Burette et al., 2015; Jiang et al., 2024). Despite these links to neurological phenotypes and the well-established role of the Wnt pathway in multiple aspects of neural development (Alkailani et al., 2022), the function of *TNIK* in NPCs, including whether it activates the Wnt signaling pathway, is unknown.

Wnt reporter assays demonstrate that TNIK can modulate Wnt signaling in NPCs. In both human and chimpanzee NPCs, inhibition of TNIK decreased Wnt reporter activity (*p* = 0.021) with no effect on control reporter activity (*p* = 0.987) (Fig. 3C; Fig. S4C). This raises the possibility that decreased RE-*TNIK* activity in human NPCs may attenuate Wnt signaling in promoting neuronal differentiation in specific contexts (Hirabayashi et al., 2004) or in generating deep-layer neurons at the expense of upper-layer neurons (Wrobel et al., 2007), which are disproportionately expanded in the human neocortex.

Further transcriptomic analyses of CRISPRi targeting RE*TNIK* in NPCs revealed genome-wide changes in the expression of genes involved in DNA replication. RNA-seq analysis of NPCs infected with either gRNAs targeting RE-*TNIK* or NTC gRNAs identified 244 differentially expressed genes in NPCs differentiated from two chimpanzee iPSC lines and 928 differentially expressed genes in NPCs differentiated from two human iPSC lines (Fig. 3D; Table S4). The weaker effect in chimpanzee NPCs is consistent with CRISPRi resulting in a 13.7% decrease in *TNIK* expression in chimpanzee NPCs compared to a 42.5% decrease in human NPCs (Fig. S4D). Genome-wide changes in gene expression were correlated between human and chimpanzee NPCs (Fig. S4E), suggesting that *TNIK* function is conserved across species. In both human and chimpanzee NPCs, inhibiting RE-*TNIK* changed the expression of genes previously implicated in the Wnt signaling pathway (Fig. 3D; Table S4), including *ACBD3, ARHGEF10, CCNL1*, and *KCTD15* (Dutta and Dawid, 2010; Huang et al., 2018; O’Brien et al., 2022; Dopeso et al., 2024). Gene-set enrichment analysis found that RE-*TNIK* inhibition affected genes involved in DNA packaging, chromatin assembly, and DNA replication-dependent nucleosome assembly in both human and chimpanzee NPCs (Fig. 3E; Table S4). These results suggest that decreased TNIK expression in human compared to chimpanzee NPCs may modulate the Wnt pathway and DNA replication, potentially affecting Wnt functions related to neuronal differentiation.

### 14 cis-regulated TFs are enriched at trans-regulated open chromatin regions in NPCs

The human-chimpanzee tetraploid system uniquely enables the identification and analysis of *cis* and *trans* gene regulatory changes within the same cell type. To determine whether analyzing *cis* and *trans* changes at the same time can identify key *cis* changes with cascading *trans* effects, we searched for *cis*regulated expression changes in transcriptional effectors that may result in widespread *trans* regulatory effects at their target sites. We focused on identifying *cis*-regulated TFs whose binding motifs are enriched at *trans*-regulated open chromatin regions, since these would drive species-differential TF availability that might affect regulatory activity at existing binding sites genome-wide (Fig. 4A). At *trans*-regulated open chromatin regions, TF motif analyses identified 282 TFs whose motifs are enriched at human-biased regions and 112 TFs whose motifs are enriched at chimpanzee-biased regions (Fig. 4B; Table S5). The greater number of human-biased TFs reflects motifs that are recognized by a substantial number of TFs; for instance, the AP-1 motif alone accounts for 42 of the human-biased TFs.

**Figure 4:**
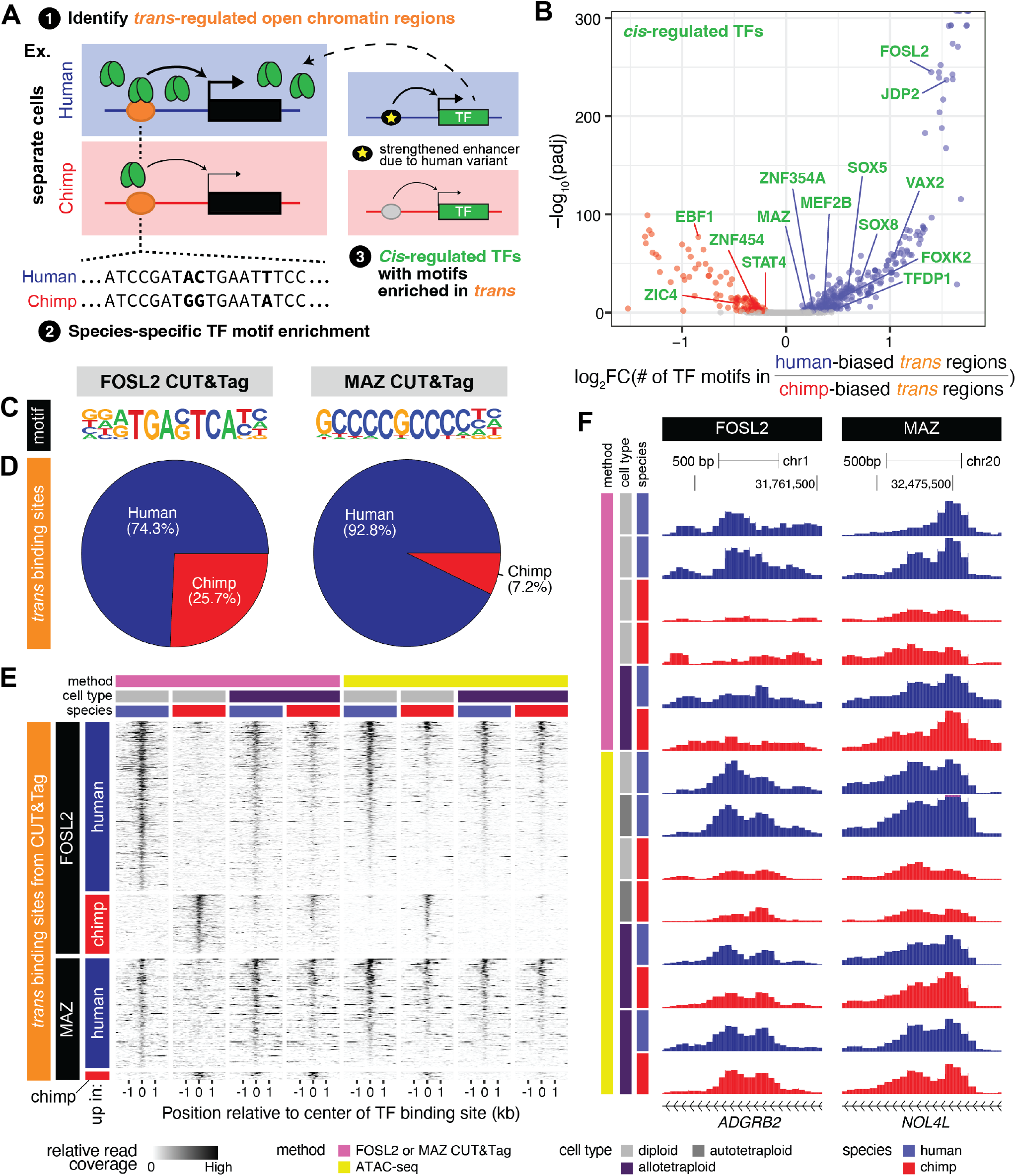
14 *cis*-regulated TFs are enriched at *trans*-regulated open chromatin regions in NPCs. (A) Workflow for identifying *cis*-regulated TFs at *trans*-regulated open chromatin regions. (B) TF motifs enriched in *trans*-regulated open chromatin regions. *Cis*-regulated TFs are labeled in green. (C) *De novo* motif discovery of binding sites from FOSL2 and MAZ CUT&Tag recovers known FOSL2 and MAZ motifs. (D) Proportion of *trans*-regulated FOSL2 and MAZ binding sites that are human-biased or chimpanzee-biased. (E) Heatmap of relative read coverage from CUT&Tag and ATAC-seq for representative human, chimpanzee, and human-chimpanzee NPCs at *trans*-regulated FOSL2 and MAZ binding sites. (F) Examples of *trans*-regulated genomic regions with stronger FOSL2 (left) or MAZ (right) binding in human NPCs compared to chimpanzee NPCs. Note that TF binding and chromatin accessibility is similar between human and chimpanzee alleles in allotetraploids, indicating that these regions are *trans*-regulated. See also Fig. S5, Table S5, and Materials and Methods.

Of the species-biased TFs, 14 are *cis*-regulated TFs (Fig. 4B), suggesting that changes in the levels of these TFs between human and chimpanzee NPCs resulted in differences in chromatin accessibility at their target sites (Fig. 4A). Ten of these *cis*-regulated TFs have motifs that are enriched at human-biased, *trans*-regulated open chromatin regions, including TFDP1, an oncogene that promotes cell cycle gene expression (Yasui et al., 2003), and SOX5, which regulates neurogenesis and neural differentiation (Martinez-Morales et al., 2010). The remaining 4 *cis*-regulated TFs have motifs that are enriched at chimpanzee-biased, *trans*-regulated open chromatin regions and include EBF1. EBF TF motifs are also enriched at chimpanzee-biased, *cis*-regulated open chromatin regions (Fig. 1F) and have been shown to be required for the initiation of neuronal differentiation (Garcia-Dominguez et al., 2003). The established functions of these TFs are consistent with an increased propensity toward self-renewal in human compared to chimpanzee NPCs. Other *cis*-regulated TFs enriched at *trans*-regulated open chromatin regions have unknown functions in NPCs, including two human-biased TFs: FOSL2, which binds the AP-1 motif, and MAZ, a Myc-associated zinc finger protein that acts as a transcriptional activator in diverse cell types (Liu et al., 2016; Maity et al., 2018) (Fig. 4B; Fig. S5A). Increased *FOSL2* and *MAZ* expression is likely derived in humans, as both genes are also more highly expressed in the developing brain in humans compared to macaques (Fig. S5B) (Zhu et al., 2018). In addition, *FOSL2* and *MAZ* are both highly intolerant to heterozygous loss-of-function mutations in humans (pLI = 1) (Lek et al., 2016), and mutations in *FOSL2* cause abnormal cognitive as well as craniofacial development (Cospain et al., 2022), indicating that quantitative changes in gene activity may have neurological consequences.

We used CUT&Tag to identify genomic binding sites for FOSL2 and MAZ in human and chimpanzee NPCs. CUT&Tag recovered the expected motifs (Fig. 4C) and confirmed the TF motif analyses that most of the *trans*-regulated binding sites for these TFs are human-biased (Fig. 4D; Table S5; Materials and Methods). FOSL2 and MAZ binding were also enriched at predicted binding sites from the ATAC-seq data (Fig. S5C). The *trans*-regulated TF binding sites that we identified are differential between human and chimpanzee NPCs, but are indistinguishable in allotetraploid NPCs (Fig. 4E, F), indicating that changes in TF binding reflect differences in TF availability between the human and chimpanzee *trans* environments rather than *cis*-acting sequence variants between humans and chimpanzees. The same pattern is observed for chromatin accessibility, although the degree of change is less dramatic (Fig. 4E, F), suggesting that additional *trans*-acting factors, such as other TFs or chromatin remodelers, may also affect chromatin accessibility at these loci. *Trans*-regulated binding sites include a FOSL2 site near the human-biased, *trans*-regulated gene *NKAIN1* and a MAZ site near the human-biased, *trans*-regulated genes *NOL4L* and *TPX2* (Fig. 4F), which have all been previously found to promote proliferation in multiple contexts (Neumayer et al., 2014; Yu et al., 2020; Koike et al., 2022; Wang et al., 2024). Taken together, these results demonstrate that examining paired RNA-seq and ATAC-seq in the human-chimpanzee tetraploid system is a powerful approach to identify *cis*-regulated TFs with genome-wide effects on *trans*-regulated chromatin accessibility.

### Increased FOSL2 and MAZ expression in human NPCs may affect neurogenesis

The 620 human-biased, *trans*-regulated FOSL2 binding sites are enriched near genes that regulate neural precursor cell proliferation and neuronal stem cell population maintenance, whereas the 215 chimpanzee-biased sites are enriched near genes involved in axon development (Fig. 5A; Table S5). This suggests that increased *FOSL2* expression in human NPCs may preferentially increase the expression of genes that promote NPC self-renewal but not genes that promote neuronal differentiation. These gene regulatory signatures were confirmed by RNA-seq on NPCs differentiated from two human iPSC lines that were infected with gRNAs targeting the *FOSL2* promoter or NTC gRNAs (Fig. S5D; Table S6). *FOSL2* knock-down activated gene sets involved in forebrain neuron differentiation and axon development (Fig. 5B), with six genes reaching individual genome-wide significance (Fig. S5E). When genes were ranked by fold change, there was a significant enrichment of FOSL2 binding sites near genes with decreased expression (*p* = 0.021; Materials and Methods), and this enrichment was most pronounced for *trans*-regulated FOSL2 binding sites (*p* = 0.015), indicating that decreased FOSL2 binding at identified binding sites affects nearby gene expression. These results suggest that *FOSL2* knockdown may bias NPCs towards terminal differentiation into neurons through small effect size changes on many genes within relevant pathways.

**Figure 5:**
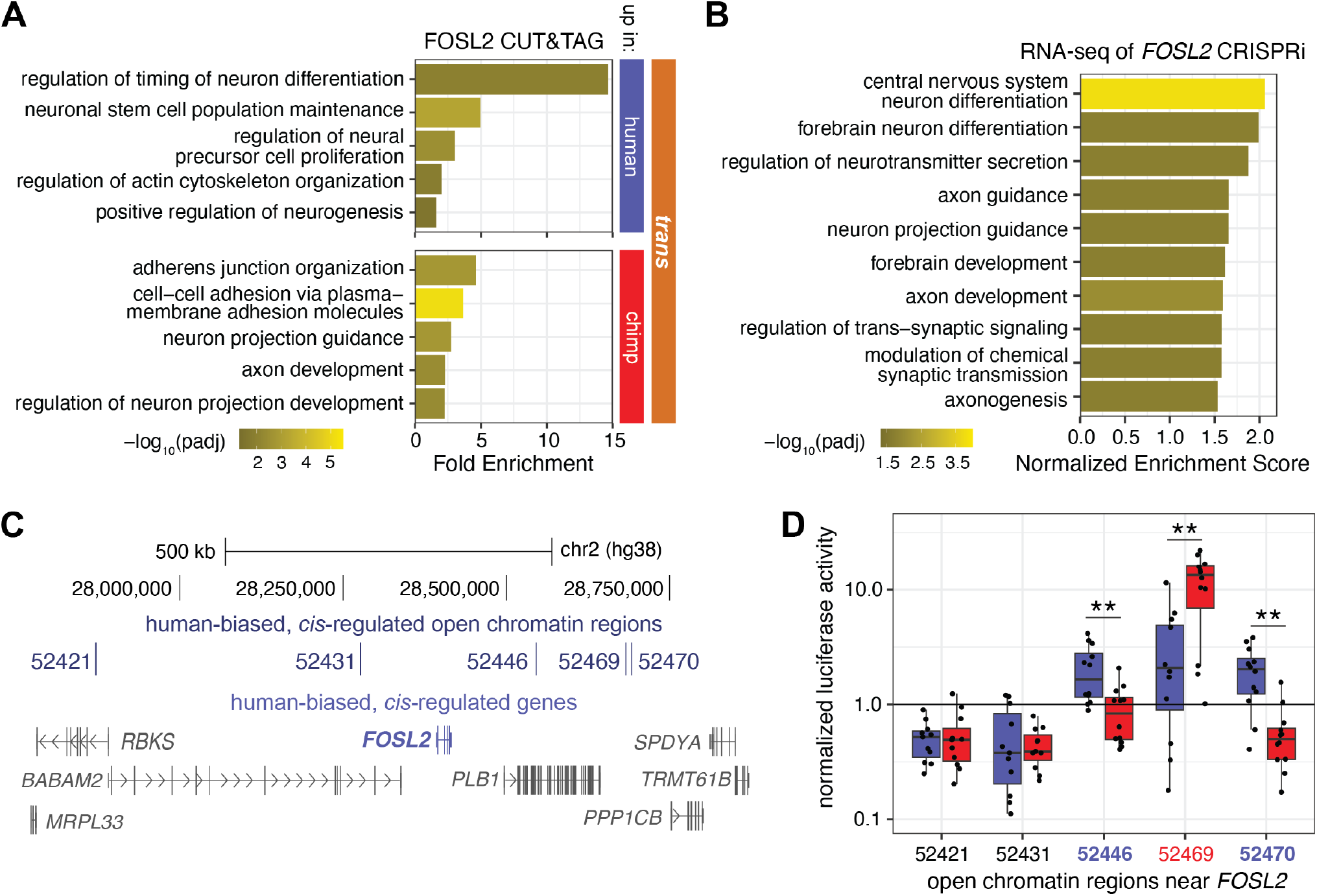
Increased *FOSL2* expression in human NPCs may affect gene programs related to neurogenesis. (A) GREAT enrichments of human-biased and chimpanzee-biased *trans*-regulated FOSL2 binding sites. (B) Gene-set enrichment analysis of RNA-seq of human NPCs infected with gRNAs targeting the *FOSL2* promoter compared to those infected with NTC gRNAs. (C) *FOSL2* is the only human-biased, *cis*-regulated gene in NPCs in its TAD. This TAD contains five human-biased, *cis*-regulated open chromatin regions. (D) Luciferase activity of the human and chimpanzee sequences of the five human-biased, *cis*-regulated open chromatin regions flanking *FOSL2*. **: *pad j* < 0.01. See also Fig. S5, Tables S5 and S6, and Materials and Methods.

Similarly, CRISPRi targeting of the *MAZ* promoter with three different gRNAs in human NPCs activated gene sets involved in neuron maturation and ion channel activity, whereas it repressed genes involved in the DNA packaging complex and oxidoreductase activity, with 11 genes reaching individual genome-wide significance (Fig. S5F-H; Table S6). Intriguingly, the expression of oxidative stress pathways is required to maintain NPCs in a state of self-renewal (Chui et al., 2020). This indicates that like *FOSL2* knockdown, *MAZ* knockdown may bias human NPCs towards terminal differentiation into neurons, albeit through distinct pathways. Taken together, these results suggest that increased *FOSL2* and *MAZ* expression in human compared to chimpanzee NPCs may promote NPC self-renewal through subtle, genome-wide shifts in expression of many target genes, demonstrating how analyzing *cis* and *trans* changes together can identify individual *cis* expression changes in key TFs with cascading, genome-wide *trans* effects.

### Cis-regulated enhancers near FOSL2 and MAZ may regulate their expression

To identify *cis*-acting sequence variation that might underlie increased expression of *FOSL2* and *MAZ* in human NPCs, we examined the *FOSL2* and *MAZ* genomic loci for human-biased, *cis*-regulated open chromatin regions. Within its TAD (Won et al., 2016; Schmitt et al., 2016; Bonev et al., 2017), *FOSL2* is the only human-biased, *cis*-regulated gene and is flanked by five human-biased, *cis*-regulated open chromatin regions (Fig. 5C). Of the five human-biased, *cis*-regulated open chromatin regions near *FOSL2*, two acted as enhancers and had significantly stronger enhancer activity from the human sequence compared to the chimpanzee sequence in NPCs by luciferase reporter assays (Fig. 5D). For the TAD containing *MAZ*, there are three human-biased, *cis*-regulated open chromatin regions, as well as other *cis*-regulated genes (Fig. S5I). Of the three human-biased, *cis*-regulated open chromatin regions near *MAZ*, one acted as an enhancer in luciferase assays and had significantly stronger enhancer activity from the human compared to the chimpanzee sequence in NPCs (Fig. S5J). These experiments nominate three potential regulatory elements, all of which contain derived human sequence variants, that may drive increased *FOSL2* or *MAZ* expression in human NPCs. Altogether, these results demonstrate how simultaneous analysis of *cis* and *trans* changes can identify species-specific gene regulatory networks that begin with *cis*-acting human-chimpanzee sequence variation affecting the expression of a nearby gene to result in cascading gene expression changes at downstream target loci in *trans*.

### Cis changes are more consistent between NPCs and excitatory neurons than trans changes

Non-coding gene regulatory variants are thought to be drivers of evolutionary change because they can avoid pleiotropic effects by restricting gene expression changes to specific cell types and developmental stages (Carroll, 2008). To examine gene regulatory changes across cell types, we compared NPCs with excitatory neurons, major building blocks of neural circuitry that result from terminal differentiation of NPCs *in vivo* and that have a prolonged maturation period in humans (Vanderhaeghen and Polleux, 2023). Our companion paper (Carter et al., 2025) differentiated multiple human and chimpanzee diploids, autotetraploids, and allotetraploid stem cell lines into cortical-like, patterned, induced excitatory neurons (piNs) using NGN2 induction combined with dual SMAD inhibition (Nehme et al., 2018). RNA-seq and ATAC-seq were performed after 28 days of differentiation.

Comparison of gene expression and chromatin accessibility between NPCs and piNs identified greater numbers of *cis* changes than *trans* changes shared between these two cell types. There were 799 genes that were *cis*-regulated in both NPCs and piNs (37% of *cis*-regulated genes in NPCs; 45% of *cis*-regulated genes in piNs), compared to only 151 genes that were *trans*-regulated in both cell types (10% of *trans*-regulated genes in NPCs; 17% of *trans*-regulated genes in piNs) (Fig. 6A). >99% of *cis*-regulated gene expression changes found in both NPCs and piNs had the same direction of effect, whereas 75% of *trans* changes had the same direction of effect (Fig. S6A). In contrast, chromatin accessibility was far more cell type-specific as expected (Corces et al., 2016), with the vast majority of open chromatin regions being unique to either NPCs or piNs. However, concordance across cell types was still higher for *cis* compared to *trans* changes in chromatin accessibility: 13% of *cis*-regulated open chromatin regions in NPCs were also *cis*-regulated in piNs, whereas only 2% of *trans*-regulated open chromatin regions in NPCs were also *trans*-regulated in piNs (Fig. 6A; Fig. S6A). Consistent with our findings in NPCs, there was an enrichment for HAQERs, but not for HARs, at *cis*-regulated open chromatin regions in piNs (Fig. S6B). Thus, although species-differential open chromatin regions were largely disjoint between NPCs and piNs, this suggests that the likely regulatory functions of HARs and HAQERs are consistent across cell types, illustrating how distinguishing both *cis* and *trans* changes can provide new insights into the functions of gene regulatory elements across cell types.

**Figure 6:**
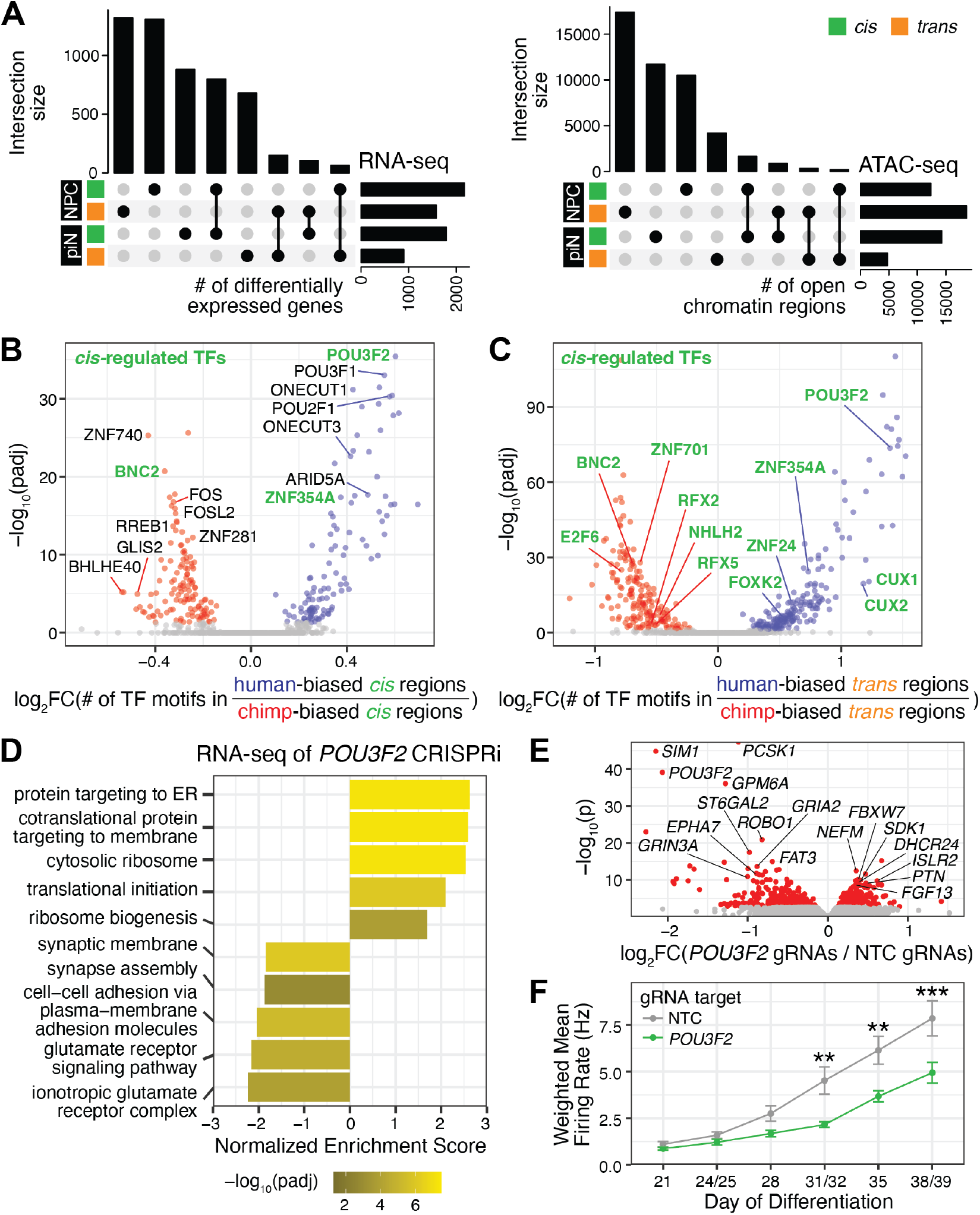
POU3F2 is a *cis*-regulated TF in piNs that affects synaptic gene expression and neuronal firing. (A) Upset plots comparing RNA-seq (left) and ATAC-seq (right) between NPCs and piNs. (B) TF motifs enriched in *cis*-regulated open chromatin regions. (C) TF motifs enriched in *trans*-regulated open chromatin regions. *Cis*-regulated TFs are labeled in green. (D) Gene-set enrichment analysis of RNA-seq of human piNs that were infected with gRNAs targeting the *POU3F2* promoter compared to those infected with NTC gRNAs. (E) Volcano plot of differential gene expression between human piNs that were infected with gRNAs targeting the *POU3F2* promoter or with NTC gRNAs. Genes significant at 5% FDR are in red. (F) Weighted mean firing rate of MEA recordings of piNs differentiated from the human line NCRM5 that were infected with gRNAs targeting the *POU3F2* promoter (green) or with NTC gRNAs (gray). Mean ± SEM is plotted. **: *pad j* < 0.01; ***: *pad j* < 0.001. See also Fig. S6, Tables S7 and S8, and Materials and Methods.

### POU3F2 is a cis-regulated TF in piNs that affects synaptic gene programs and neuronal firing

Motif analyses of open chromatin regions in piNs identified TFs that may affect chromatin accessibility (Fig. 6B, C; Table S7). Motifs for POU and ONECUT TFs, which are known to affect neuronal differentiation and maturation (van der Raadt et al., 2019; Ding et al., 2021), are enriched at human-biased, *cis*-regulated open chromatin regions. In contrast, motifs for TFs like RREB1, which regulates dendritic development (Griffin et al., 2024), are enriched at chimpanzee-biased, *cis*-regulated open chromatin regions, potentially representing a molecular signature of faster neuronal maturation. At *trans*-regulated open chromatin regions, we identified motifs for six *cis*-regulated TFs enriched at human-biased sites and six *cis*-regulated TFs enriched at chimpanzee-biased sites. Notably, motifs for three *cis*-regulated TFs, POU3F2, ZNF354A, and BNC3, are enriched at both *cis*- and *trans*-regulated open chromatin regions, suggesting that changes in the levels of these TFs may mediate downstream *trans* changes at their existing binding sites and that these effects may be further amplified by *cis*-acting sequence variants that improve TF motif strength. We further investigated POU3F2, a TF that is expressed throughout neural development and plays critical roles in neurogenesis and neuron differentiation (Zhu et al., 2022; Lu et al., 2023). *POU3F2* is differentially expressed between humans and chimpanzees in piNs but not in NPCs (Table S1) (Carter et al., 2025). In human piNs, CRISPRi targeting of the *POU3F2* promoter with three different gRNAs activated gene sets involved in protein targeting and translation while inhibiting gene sets involved in synapse assembly and glutamate signaling (Fig. 6D; Fig. S6C, D; Table S8). Genome-wide, there were 672 differentially expressed genes by RNA-seq (Fig. 6E). There was no discernable difference in marker gene expression, suggesting that *POU3F2* knockdown did not affect piN differentiation (Fig. S6D). Given that *POU3F2* knockdown affected the expression of synapse-related genes, we tested whether *POU3F2* may affect neuronal firing.

Indeed, multi-electrode array (MEA) recordings of neuronal firing rates from day 21 to day 39 of differentiation revealed that *POU3F2* knockdown significantly decreased firing rates in piNs differentiated from two human diploid lines (Fig. 6F; Fig. S6E). Taken together, knockdown of *POU3F2*, a *cis*-regulated TF enriched at human-biased open chromatin regions in piNs, inhibited the expression of synapse-associated genes and decreased neuronal firing rates. Given that *POU3F2* is more highly expressed in human compared to chimpanzee piNs, this suggests that an evolutionary increase in *POU3F2* expression in human piNs affected *cis*- and *trans*-regulated chromatin accessibility, activated synapse-associated genes, and modulated neuronal firing rates. These findings demonstrate how the ability to distinguish *cis* and *trans* changes using the human-chimpanzee tetraploid system can identify *cis*-regulated transcriptional effectors with genome-wide effects on *trans*-regulated gene expression, not only in NPCs, but also in excitatory neurons.

## Discussion

By distinguishing *cis*- and *trans*-regulated gene expression and chromatin accessibility changes between human and chimpanzee NPCs and piNs, the human-chimpanzee tetraploid system allowed us to identify a wide range of genes and regulatory regions with significant, species-specific regulatory effects during the process of neural proliferation and maturation. The identification and characterization of RE-*TNIK* demonstrates how the tetraploid system can directly identify *cis*-acting sequences with species-specific effects in an unbiased fashion, orthogonal and complementary to the identification of candidate elements by phylogenetic comparison of DNA sequences (e.g. HARs and HAQERs). We also established a second, independent approach using the human-chimpanzee tetraploid system where we analyzed both *cis* and *trans* changes to identify *cis*-regulated transcriptional effectors, such as FOSL2, MAZ, and POU3F2, that have genome-wide *trans* effects on TF binding, gene expression, and neural properties. Together, these results identify a host of specific genetic changes that likely contribute to changes in NPC proliferation and neuronal maturation in humans and provide a roadmap for the use of human-chimpanzee tetraploids to characterize human evolution across diverse cell types.

Most of the >20 million variants between humans and chimpanzees are likely neutrally evolving with little to no functional effect, making the identification of critical functional variants enormously challenging (Fair and Pollen, 2023). To begin to identify which of the >12,000 *cis*-regulated open chromatin regions in NPCs are necessary for species-differential gene expression, we tested 106 of these regions in a CRISPRi screen and identified 7 hits, in line with the 0.3 ™ 10% of target regions that affected gene expression in previously published CRISPRi screens (Gasperini et al., 2019; Fair et al., 2023; Geller et al., 2024). In part, these modest hit rates reflect enhancer redundancy; we observed an increased proportion of human-biased open chromatin regions near human-biased genes and vice versa, and experimentally verified two species-differential enhancers that regulate *TNIK* expression. They likely also reflect the fact that most variants have negligible functional effects, suggesting that CRISPRi screens may be an effective, high-throughput approach to quickly identify the small percentage of variants necessary for gene expression. Modifying the selection of *cis*-regulated open chromatin regions to test — by prioritizing genomic loci with fewer *cis*-regulated open chromatin regions to avoid enhancer redundancy, or by targeting novel human regulatory elements like HAQERs by not requiring sequence conservation in non-human primates — has the potential to uncover additional *cis*-regulated open chromatin regions necessary for species-differential gene expression in NPCs.

Our combination of genome-wide surveys and functional CRISPRi testing strongly implicate Wnt signaling in NPC evolution. Three of the hits, including RE-*TNIK*, regulated genes previously linked to Wnt signaling (Shitashige et al., 2010; Coba et al., 2012; Durak et al., 2015; Lee et al., 2017; Bahn and Ko, 2023), and we found that RE-*TNIK* modulates cell cycle gene expression. In different contexts, activating the Wnt pathway in neural progenitors can promote neurogenesis (Chenn and Walsh, 2002), neuronal differentiation (Hirabayashi et al., 2004), or the generation deep-layer neurons at the expense of upper-layer neurons (Wrobel et al., 2007). These seemingly conflicting functions may explain why we identified known Wnt signaling activators with both increased (e.g. *ANK3*) and decreased (e.g. *TNIK*) expression in human NPCs. Together with the prior identification of a HAR that regulates the Wnt receptor *FZD8* (Boyd et al., 2015; Liu et al., 2024), these results suggest that the Wnt signaling pathway was a major site of evolutionary tinkering to regulate human neurodevelopment. The ability to dissect different types of gene regulation can enable functional insights into genomic regions with signatures of selection in the human lineage, a long-standing interest in the field. We find that HARs, a set of conserved regulatory elements that change in humans (Girskis et al., 2021; Keough et al., 2023; Bi et al., 2023), do not show statistically significant enrichment for *cis*-regulated open chromatin regions in either NPCs or piNs, lending credence to recent findings that conserved genomic elements that uniquely change in humans often contain sequence variants with opposing effects on enhancer activity (Whalen et al., 2023) or may merely compensate for changes in the *trans* environment (Fair et al., 2023). In contrast, HAQERs, a set likely composed of novel human regulatory elements (Mangan et al., 2022), are enriched for *cis*-regulated open chromatin regions in both NPCs and piNs. This may suggest that HARs, defined by their strong conservation in non-humans, mainly comprise elements under strong negative selective pressure to maintain an existing regulatory element, such that sequence changes in HARs are more likely to result in non-directional or compensatory changes, whereas novel regulatory elements like HAQERs are less constrained and more likely to have *cis* effects. Thus, an effective strategy to enrich for genetic changes with strong evolutionary effects may be to intersect *cis*-regulated regions identified here by the human-chimpanzee tetraploid system with previously identified genomic regions with signatures of selection in the human lineage.

Distinguishing and examining both *cis* and *trans* changes with the human-chimpanzee tetraploid system provides a unique opportunity to identify species-specific changes in experimentally grounded gene regulatory networks. Here, we identified TFs with *cis*-regulated gene expression differences, where changes in TF availability between humans and chimpanzees result in cascading gene regulatory changes in *trans*. We showed that FOSL2 and MAZ are *cis*-regulated TFs with hundreds of genome-wide *trans* effects on chromatin accessibility and TF binding in NPCs. When either gene was depleted, we observed clear transcriptomic shifts toward gene programs involved in neuron differentiation, suggesting that increased expression of these TFs in human NPCs may have promoted NPC self-renewal during mitosis. We further identified POU3F2 in piNs as a *cis*-regulated TF whose targets are enriched at human-biased open chromatin regions with strong *trans* effects on synaptic gene expression and neural firing rates. Future identification of the exact *cis*-acting sequence variants that drive differential TF expression and re-creation of those variants in their endogenous context will more precisely determine their evolutionary consequences. In contrast to prior studies that focused solely on *cis* changes when using the human-chimpanzee tetraploid system (Agoglia et al., 2021; Gokhman et al., 2021; Wang et al., 2023; Carter et al., 2025), our results illustrate how simultaneously analyzing *cis* and *trans* changes can uncover key transcriptional effectors with substantial downstream effects.

This study demonstrates multiple approaches to identify key evolutionary genetic changes using the human-chimpanzee tetraploid system in neural progenitors and neurons, and illustrates how distinguishing *cis* and *trans* regulation can enrich our functional understanding of genomic regions identified by orthogonal approaches, such as through DNA sequence comparisons (McLean et al., 2011; Girskis et al., 2021; Mangan et al., 2022; Keough et al., 2023; Xue et al., 2023; Bi et al., 2023). The approaches pioneered here are agnostic to existing domain knowledge and can be readily applied to any cell type or state, including ones where relatively little is known about their molecular landscape, providing critical entry points for understanding human-specific gene regulatory changes in diverse cellular contexts. Ultimately, further examination of the genetic changes nominated by this approach in single locus models will provide a detailed understanding of the key genomic changes contributing to human evolution.

## Acknowledgements

We thank members of the Walsh and Greenberg labs for useful discussions and the BCH FACS core, the BCH Human Neuron Core, and the BCH IDDRC Molecular Genetics Core Facility for equipment use and assistance. J.H.T.S. is supported by the Y. Eva Tan Fellowship from the Tan Yang Autism Center at Harvard University, the Howard Hughes Medical Institute Fellowship from the Helen Hay Whitney Foundation, and grant no. K99MH136290 from NIMH/NIH. A.C.C. is supported by the Hanna H. Gray Fellowship from the Howard Hughes Medical Institute. C.A.W. and M.E.G. are supported by the Allen Discovery Center program, a Paul G. Allen Frontiers Group advised program of the Paul G. Allen Family Foundation, and the Hock E. Tan and K. Lisa Yang Center for Autism Research at Harvard University. C.A.W. is supported by the Templeton Foundation. C.A.W. and D.M.K. are Investigators of the Howard Hughes Medical Institute. This study makes use of data generated by the DECIPHER community. A full list of centres who contributed to the generation of the data is available from https://deciphergenomics.org/about/stats and via email from. DECIPHER is hosted by EMBL-EBI and funding for the DECIPHER project was provided by the Wellcome Trust [grant number WT223718/Z/21/Z]. The content of this paper is solely the responsibility of the authors and does not necessarily represent the official views of the NIH.

## Author Contributions

**Janet H.T. Song**: Conceptualization, Methodology, Formal Analysis, Investigation, Writing - Original Draft, Writing - Review & Editing, Visualization. **Ava C. Carter**: Conceptualization, Methodology, Formal Analysis, Investigation, Writing

- Original Draft, Writing - Review & Editing, Visualization. **Evan M. Bushinsky**: Investigation. **Samantha G. Beck**: Investigation. **Jillian Petrocelli**: Investigation. **Gabriel Koreman**: Methodology, Formal Analysis, Investigation, Writing - Original Draft, Writing - Review & Editing. **Juliana Babu**: Methodology, Investigation. **David M. Kingsley**: Resources, Writing - Review & Editing. **Michael E. Greenberg**: Conceptualization, Writing - Original Draft, Writing - Review & Editing. **Christopher A. Walsh**: Conceptualization, Writing - Original Draft, Writing - Review & Editing.

## Materials and Methods

### Cell lines

Information about each cell line used in this study is listed in Table S9 (Gallego Romero et al., 2015; Tian et al., 2019; Song et al., 2021; She et al., 2023). All cell lines were maintained in a 5% CO_2_ incubator at 37°C.

### Cell culture

Stem cell lines (Table S9) were routinely propagated feeder-free in mTeSR Plus (STEMCELL Technologies, cat #100- 0276). Stem cell lines were differentiated into neural progenitor cells (NPCs) using the StemDiff SMADi Neural Induction Kit (STEMCELL Technologies, cat #08582) as per the manufacturer’s instructions. Briefly, 2*x*10^6^ stem cells were plated per well of a 6-well plate in 2ml of Neural Induction Media + 1µM thiazovivin (Tocris, cat #3845) as D0. Media was changed to fresh Neural Induction Media without thiazovivin each day. Cells were split on D7 using Accutase (Millipore, cat #SCR005) and 2*x*10^6^ stem cells were plated per well of a 6 well plate in 2ml of Neural Induction Media + 1µM thiazovivin. Media was changed to fresh Neural Induction Media without thiazovivin each day. Cells were typically harvested at D14. Plates for stem cell maintenance and for NPC differentiation were precoated with geltrex (Gibco, cat #A1413202). For luciferase assays, 2*x*10^6^ cells were frozen per vial at D7 and thawed for use in subsequent experiments.

Stem cells were differentiated into cortical-like, patterned, induced neurons (piNs) as previously described (Nehme et al., 2018) with minor modifications as detailed in (Carter et al., 2025).

### Immunofluorescence

NPCs were split at D12-13 and 2*x*10^4 ™^ 1*x*10^5^ cells were plated in 1 chamber of a 4-chamber slide (Thermo Scientific, cat #155382). Two days after plating, NPCs were rinsed in 1X PBS and fixed in 4% PFA for 15 minutes at room temperature. NPCs were then washed for 3 × 5min in 1X PBS, incubated in blocking solution (8% donkey serum, 0.3% BSA, 0.3% Triton X-100, 0.02% sodium azide in 1X PBS) for 1h at room temperature, and incubated in primary antibodies diluted in blocking solution overnight at 4°C. The next day, slides were washed for 3 × 5min in 1X PBS, incubated in donkey secondary antibodies (Invitrogen) diluted in blocking solution for 2-3h at room temperature, stained with DAPI (Invitrogen, cat #R37606), and washed for 3 × 5min in 1X PBS. Slides were imaged on the Axio Observer Z1 microscope (Zeiss) in 1X PBS. Primary antibodies: mouse anti-Nestin (Millipore, cat #MAB5326) at 1:200, rabbit anti-Pax6 (Biolegend, cat #901302) at 1:500, goat anti-Sox2 (R&D, cat #AF2018) at 1:200.

### Generation of stem cell lines for CRISPR inhibition

We inserted dCas9-BFP-KRAB into the CLYBL locus in two chimpanzee iPSC lines (C2, C3) and one human iPSC line (H2) (Table S9). A sgRNA targeting the CLYBL locus, CLYBL-gRNA (Table S9), was *in vitro* transcribed as previously described (Wucherpfennig et al., 2019). 1µL of 40µM Cas9-NLS purified protein (QB3, UC Berkeley) was complexed with 2-3µg of sgRNA for 5 minutes at room temperature. This complex and 5-10µg of the donor plasmid pC13N-dCas9-BFPKRAB (Tian et al., 2019) (Addgene, cat #127968) were nucleofected into 1*x*10^6^ cells with the Neon transfection System 100µl Kit (Thermo Fisher, cat #MPK10096) with the following conditions: pulse voltage = 1400, pulse width = 20, pulse number = 1 for C2 and H2; pulse voltage = 1100, pulse width = 30, pulse number = 1 for C3.

Following nucleofection, cells were plated in a 10cm plate in mTeSR Plus + 1µM thiazovivin. 2-3 days later, cells containing the donor plasmid were selected using 1mg/ml G418 (Thermo Fisher, cat #10131035) in mTeSR Plus for 3-5 days. Cells were then allowed to recover for 7-10 days and form colonies. Colonies were picked using the EVOS M5000 microscope (Thermo Fisher) into a 96-well plate. When colonies were confluent, they were split 1:5 into a new 96-well plate. The remaining cells were resuspended in TE buffer, heated at 95°C for 10 min to lyse the cells, incubated at 55°C for 1h with Proteinase K (Thermo Fisher, cat #25530015), and then heated at 95°C for 10 min to denature the Proteinase K. 1/20 of each sample was used for genotyping with primers spanning the 5’ and 3’ junctions and the entire CLYBL locus (Table S9). Clones with insertions were confirmed by Sanger sequencing, and expanded. Clones expressing the inserted transgene were selected by multiple rounds of FACS for BFP expression. dCas9-KRAB expression was confirmed by qPCR (Table S9). Karyotype analysis was performed by WiCell.

### RNA isolation, qPCR, library preparation, and sequencing

NPCs at D14 or piNs at D28 were collected in TRIzol and RNA was extracted with the Direct-Zol Miniprep Kit (Zymo Research, cat #R2052) or the Direct-Zol Microprep Kit (Zymo Research, cat #R2062). qPCR was performed on the CFX384 Touch Real-Time PCR Detection System (Bio-Rad) with qPCR Brilliant II SYBR MM with ROX (Agilent, cat #600830) and primers for the normalizer gene *GAPDH* and the appropriate target gene (Table S9), after RNA was reverse transcribed using the SuperScript VILO kit (Life Technologies, cat #11755050). RNA sequencing libraries were made with the SMARTer Stranded Total RNA-Seq Kit v2 - Pico Input Mammalian kit (Takara Bio) or with the WatchMaker RNA Library Prep Kit with Polaris Depletion (WatchMaker Genomics) and the Twist UMI Adapter System - TruSeq Comp, 96 Samples Plate A (Twist Biosciences, cat #105041). Samples were sequenced on Illumina NovaSeq platforms with paired-end 150bp reads. See Tables S1, S4, S6, S8 for the number of samples and associated metadata for each sequencing experiment.

### ATAC library preparation and sequencing

At D14 of differentiation, NPCs in a 12-well plate were lyzed in 250µl of lysis buffer (10mM Tris-HCl pH7.4, 10mM NaCl, 3mM MgCl_2_, 0.1% NP-40, 0.1% Tween-20) per well and then scraped and collected into conical tubes at a final volume of 500µl lysis buffer. The cell suspension was incubated at 4°C for 10min to release nuclei and then pelleted at 830rcf for 10min. Nuclei were resuspended in 1ml lysis buffer and counted. 50,000 nuclei were then pelleted at 830rcf for 10min and resuspended in 50µl of transposition mix (1X TD buffer [10mM Tris-HCl pH7.6, 5mM MgCl_2_, 10% dimethyl formamide], 0.33X PBS, 0.001% Digitonin, 0.1% Tween-20, 100nM Tn5 transposase [made in house]). The resuspended nuclei were incubated at 37°C for 30min. DNA was then purified and eluted in 21µl H2O using the DNA Clean and Concentrator-5 Kit (Zymo Research, cat #D4014). 20µl of the eluent was PCR amplified with 1X NEBNext High Fidelity PCR Master Mix (New England Biolabs, cat #M0541L), 1.25µM Ad1, and 1.25µM Ad2 (primer sequences in Table S9) for 72°C for 5min, 98°C for 30s, and then 5 cycles of 98°C for 10s, 63°C for 30s, and 72°C for 1min. Libraries were purified using the DNA Clean and Concentrator-5 Kit and quantified with the KAPA Library Quantification Kit (Roche, cat #07960140001). Samples were sequenced on the Illumina NovaSeq 6000 with paired-end 150bp reads. See Table S2 for the number of samples and associated metadata for ATAC-seq.

### CRISPR inhibition

Stem cell lines with constitutive expression of dCas9KRAB were used in all CRISPR inhibition (CRISPRi) experiments (Table S9). gRNA sequences (Table S9) were cloned into pBA904 (Replogle et al., 2020) (Addgene, cat #122238) expressing BFP, GFP, or mCherry using BstXI (Thermo Fisher, cat #FD1024) and BlpI (Thermo Fisher, cat #FD0094). Lentivirus was generated with third generation lentiviral plasmids in 293T cells followed by ultracentrifugation, and titered in NPCs. NPCs were infected at D7 of differentiation at MOI 0.7, and infected cells were sorted by FACS for the relevant fluorophore at D14. Sorted cells were then collected in TRIzol for RNA extraction. piNs were infected and processed as described in (Carter et al., 2025).

For the experiment shown in Fig. 2F and Fig. S2E, cells were infected with two lentiviruses, one expressing GFP and the other expressing mCherry. The lentiviruses also expressed one of three NTC gRNA (NTC2, NTC3, NTC5), a gRNA targeting the *TNIK* promoter (TNIK TSS2), a gRNA targeting RE-*TNIK* (72060-2), or one of three gRNAs targeting RE2-*TNIK* (72061-0, 72061-6, 72061-7) (Table S9). Infections were performed for all combinations (2 NTC gRNAs, 1 NTC gRNA and 1 test gRNA, or 2 test gRNAs) and fluorophores were mixed between NTC and test gRNAs (ex. NTC5 and 72060-2 were used with both GFP and mCherry). GFP+, mCherry+, and GFP+/mCherry+ cells were sorted with FACS and collected in TRIzol for RNA extraction.

### CRISPR inhibition screen

The gRNA library was cloned into pBA904 (Addgene, cat #122238) as previously described: https://weissman.wi.mit.edu/resources/PooledCRISPRLibraryCloning.pdf. Briefly, an oligo pool containing each gRNA (Table S3) was synthesized by IDT, PCR amplified for 8 cycles, purified with the MinElute PCR Purification Kit (Qiagen, cat #28204), and digested with BstXI (Thermo Fisher, cat #FD1024) and BlpI (Thermo Fisher, cat #FD0094). The digested product (∼ 33bp) was gel extracted by crushing the gel through a hole in a 0.5ml nonstick tube by centrifugation, adding 200µl of water, and incubating for 1h at 70°C. The gel slurry was then filtered through a Costar Spin-X column (Millipore Sigma, cat #CLS8160-24EA) followed by ethanol precipitation. In parallel, pBA904 was digested with BstXI and BlpI and gel extracted using the QIAquick Gel Extraction Kit (Qiagen, cat #28704).

The digested gRNA pool was ligated to the digested pBA904 plasmid at a 1:2 molar ratio and transformed into 33µl of MegaX DH10B T1R Electrocomp Cells (Thermo Fisher, cat #C640003). The transformation was plated on a large tray (Fisher, cat #NC9372402) with LB and 75ng/ml carbenicillin (Fisher, cat #BP26481) and grown for 14-17h at 37°C. Bacterial colonies were then collected in LB/carbenicillin using a cell scraper and plasmid DNA was extracted using the NucleoBond Xtra Midi Plus EF kit (Macherey-Nagel, cat #740422.5). gRNA complexity was confirmed by PCR of the plasmid library with mU6-F and pBA904-TR (Table S9) followed by amplicon-EZ sequencing (GeneWiz).

Lentivirus was generated from the gRNA plasmid library with third generation lentiviral plasmids in 293T cells followed by ultracentrifugation, and titered in NPCs. C3-C and NCRM5 (Table S9) were differentiated into NPCs, and passaged and infected with the lentiviral gRNA pool at D7 at MOI 0.15. At D14, infected cells (BFP+) were sorted by FACS and 4 scRNA-seq reactions were prepared for each cell line using the Chromium Next GEM Single Cell 3’ Reagent Kit v3.1 (Dual Index) with Feature Barcoding technology for CRISPR Screening (10X Genomics, cat #PN-1000268, #PN-1000269, #PN-1000262, #PN-1000120, #PN-1000127, #PN-1000215, #PN-1000242). Libraries were sequenced on NovaSeq instruments (MedGenome). There were ∼15,000 cells/reaction for NCRM5 and ∼13,000 cells/reaction for C3-C. Gene expression libraries were sequenced to ∼40,000 reads/cell and CRISPR feature barcoding libraries were sequenced to ∼8,000 reads/cell.

### Luciferase assays

NPCs frozen at D7 were thawed and allowed to recover for 2-4 days. NPCs were split and plated at 2 ™ 4*x*10^4^ cells/well in 96 well plates in 75µl of Neural Induction Media (STEM-CELL Technologies, cat #08582) and 1µM thiazovivin. The next day, NPCs were moved to 75µl of Neural Induction Media and transfected.

For enhancer reporter assays, test genomic regions were PCR amplified and cloned into the multiple restriction site in pNL3.2 (Promega, cat #N1041). The same primers were used to PCR amplify from both human DNA (H2) and chimpanzee DNA (C2) for each test region (Table S9). For RE-*PLAAT3*, reporter plasmids were ordered from VectorBuilder in lieu of cloning. NPCs were transfected with 0.2µl Lipofectamine Stem Transfection Reagent (Thermo Fisher, cat #STEM00003), 10ng of pGL4.53 (Promega, cat #E5011), and 190ng of pNL3.2 or pNL3.2 containing a test genomic region in 10µl of Opti-MEM (Thermo Fisher, cat #31985070). 24 hours later, luciferase activity was assessed with the Nano-Glo Dual Luciferase Reporter Assay System (Promega, cat #N1620). There were three technical replicates per plasmid per experiment. The enhancer reporter assays were performed in NPCs differentiated from the stem cell lines C2, C2-C, C3-C, H2, NCRM5, and H1-ESC-CRi (Table S9).

For the Wnt reporter assays, NPCs were transfected with 0.2µl Lipofectamine Stem Transfection Reagent, 2ng of pNL1.1.TK (Promega, cat #N1501), and 198ng of TOPflash (Addgene, cat #12456) or FOPflash (Addgene, cat #12457) (Veeman et al., 2003) in 10µl Opti-MEM. The next day, 0-50µM of CHIR-99021 (MedChemExpress, cat #HY-10182) with or without 50µM of KY-05009 (MedChemExpress, cat #HY-124745) was added in 75µl of fresh Neural Induction Media. 24 hours later, luciferase activity was assessed with the Nano-Glo Dual Luciferase Reporter Assay System (Promega, cat #N1620). There were three technical replicates per condition per experiment. The Wnt reporter experiments were performed in NPCs differentiated from the stem cell lines C2, C3-C, and NCRM5 (Table S9).

### Western blotting

NPCs at D14 were scraped into RIPA buffer, incubated on ice for 20min, and centrifuged at 4°C to pellet debris. The supernatant was recovered and quantified on a spectrophotometer. 20µg per sample was denatured in NuPAGE LDS Sample buffer (Life Technologies, cat #NP0007) at 95°C for 5min and loaded in NuPage 4-12% Bis-Tris Gel (Life Technologies, cat #NP0323BOX) along with the Precision Plus Protein Dual Color Standard (Bio-Rad, cat #1610374). After running the gel, the gel was transferred to a nitrocellulose membrane (BioRad, cat #10484060) overnight at 4°C at 30V. The next day, the membrane was blocked in superblock (Life Technologies, cat #37518) for 1h at room temperature and then incubated in primary antibody overnight at 4°C. Primary antibodies: rabbit anti-TNIK at 1:1000 (GeneTek, cat #GTX13141), rabbit anti-H3 at 1:5000 (Abcam, cat #ab1791). The next day, the membrane was washed 4 × 5min in 1X TBS-T, incubated for 2h with the secondary antibody goat anti-rabbit 800 (LicorBio, cat #926-32211), washed 4 × 5min in 1X TBS-T, and imaged on the LiCor Odyssey Imager.

### CUT&Tag library preparation and sequencing

At D14, NPCs were scraped off of tissue culture plates into NE1 lysis buffer (20mM HEPES pH 7.9, 10mM KCl, 0.1% Triton X-100, 3mM MgCl_2_, 0.5mM Spermidine, 1 tablet Complete Protease Inhibitor EDTA-free per 50ml) and nutated at 4°C for 10min to isolate nuclei. Nuclei were pelleted at 4°C at 200rpm for 10min and then resuspended in 1ml WB150 (20mM HEPES pH 7.5, 150mM NaCl, 0.2% Tween-20, 1g/L BSA, 0.5mM spermidine, 10mM sodium butyrate, 1 tablet Complete Protease Inhibitor EDTA-free per 50ml) for counting. For each library, 500,000 nuclei were aliquoted into a total volume of 1ml WB150. 20µl Concanavalin A bead (Bangs Laboratories, cat # BP531) slurry was prepared per library by washing beads twice in binding buffer (20µM HEPES-KOH pH7.9, 10µM KCl, 1µM CaCl_2_, 1µM MnCl_2_) and resuspending in 10µl binding buffer. Beads were added to the nuclei suspension and incubated for 10min at room temperature. Using a magnet, the supernatant was removed, and beads were resuspended in 50µl antibody buffer (WB150 supplemented with 0.1% Triton X-100 and 2mM EDTA) with 1µl primary antibody (rabbit anti-FOSL2: Cell Signaling Technology, cat #19967; rabbit anti-MAZ: Abcam, cat #ab85725; rabbit IgG: Cell Signaling Technology, cat #2729S) for overnight incubation at 4°C.

The next day, the primary antibody solution was removed and beads were resuspended in 100µl of antibody buffer with 1.2µl of secondary antibody (goat anti-rabbit: Fisher Scientific, cat #NBP172763; goat anti-mouse: Abcam, cat #ab46540). Beads were incubated at room temperature for 1h and then washed 3X with 1ml WB150. Beads were resuspended in 100µl WB300 (20mM HEPES pH 7.5, 300mM NaCl, 0.2% Tween-20, 1g/L BSA, 0.5mM spermidine, 10mM sodium butyrate, 1 tablet Complete Protease Inhibitor EDTA-free per 50ml) containing 2.5µl proteinAG-Tn5 (Epicypher Inc, cat #15-1017) and incubated at room temperature for 1h. Then, beads were washed 3X with 1ml WB300 and incubated in tagmentation buffer (WB300 supplemented with 10mM MgCl_2_) at 50°C for 1h at 500rpm. DNA was extracted with phenol/chloroform, ethanol precipitated with 0.02mg/ml GlycoBlue (Life Technologies, cat #AM9515), and resuspended in 22µl H2O.

For library preparation, 20µl of DNA was PCR amplified with 5X KAPA Hifi Buffer, 0.3mM dNTP, 1µl KAPA HiFi Polymerase (Roche, cat #KK2101), 0.4µM Ad1, and 0.4µM Ad2 (primer sequences in Table S9) in a 50µl reaction at 72°C for 5min, 98°C for 30s, 14 cycles of 98°C for 10s, 63°C for 30s, 72°C for 1min, and 72°C for 5min. DNA was purified using double-sided size selection (0.5X, then 1.2X) with Ampure XP beads (Thermo Fisher, cat #NC0110018). Briefly, 25µl of beads were added to the PCR reaction, incubated at room temperature for 8 min, and allowed to adhere to the magnet for 8 min. The supernatant was moved to a new tube and 65µl of beads were added. After 8 min incubation at room temperature, beads were allowed to adhere to the magnet for 8 min, the supernatant was discarded, and beads were washed twice with 200µl 80% EtOH for 30s each. Beads were resuspended in 53µl 10mM Tris-HCl, pH 7.5 and placed on the magnet, and the supernatant was moved to a new tube. The supernatant was then further cleaned by adding 55µl beads, incubated at room temperature for 8 min, washed twice with 200µl 80% EtOH for 30s each, and then eluted in 20µl 10mM Tris-HCl, pH 7.5. Libraries were quantified with the KAPA Library Quantification Kit (Roche, cat #07960140001) and sequenced on the Illumina NovaSeq X with paired-end 150bp reads. See Table S5 for the number of samples and associated metadata.

### Multi-electrode array (MEA) recordings

MEA recordings were performed as described in (Carter et al., 2025). Briefly, human iPSC-derived astrocytes (nCardia, cat #K0101) were plated in a 48-well CytoView MEA plate (Axion Biosystems, cat #M768-tMEA-48B) for 3 days. At day 7 of piN differentiation, neurons were dissociated and mixed with astrocytes. A mix of 60,000 neurons and 9,000 astrocytes were plated in each well in complete neurobasal (cNB) media (Neurobasal [Thermo Fisher, cat #21103049], 1% MEM-NEAA, 1% Penicillin/Streptomycin, 1% GlutaMAX [Life Technologies, cat #35050061], 1X N2 Supplement-B [STEMCELL Technologies, cat #07156], 1X B27 without vitamin A [Life Technologies, cat #12587010], 0.1% Mouse laminin [Life Technologies, cat #23017015], 20ng/ml rhBDNF [Peprotech, cat #450-02], 10ng/ml rhGDNF [Peprotech, cat #450-10], 1µM L-Ascorbic Acid [Sigma, cat #A4544], 20µM Dibutyryl cAMP [Sigma, cat #D0260]) supplemented with 10µM ROCK-inhibitor Y-27632 (STEMCELL Technologies, cat #72304). Half media changes were performed every 3 days with cNB media until day 14, after which media was replaced with complete BrainPhys media (BrainPhys Neuronal Media [STEMCELL Technologies, cat #05790], 1% Penicillin/Streptomycin, 1X N2 Supplement-A [STEMCELL Technologies, cat #07152], NeuroCult SM1 Neuronal Supplement without Vitamin A [STEMCELL Technologies, cat #05731], 0.1% Mouse laminin [Life Technologies, cat #23017015], 20ng/ml rhBDNF [Peprotech, cat #450-02], 10ng/ml rhGDNF [Peprotech, cat #450-10], 1µM L-Ascorbic Acid [Sigma, cat #A4544], 20µM Dibutyryl cAMP [Sigma, cat #D0260]). Half media changes were completed every ∼ 3 days and always immediately after recording.

Recordings were performed as described in (Carter et al., 2025) using the Axion Maestro Pro (Maestro-625) recording system.

### Differential expression (DE) and allele-specific expression (ASE) analyses of diploids, autotetraploids, and allotetraploids

RNA-seq data from the human and chimpanzee diploids, autotetraploids, and allotetraploids (Fig. 1) was analyzed as previously described (Song et al., 2021). Briefly, sequencing reads were trimmed for adapter sequences and assessed for sequencing quality using trimgalore v0.6.6 with cutadapt v2.5 (Martin, 2011) and fastqc v0.11.5 (Andrews et al., 2010). Reads were aligned using STAR v2.7.9a with two-pass mapping (Dobin et al., 2013) to a composite human-chimpanzee genome (hg38 and panTro6). The number of uniquely-mapped reads (*MAPQ* = 255) that overlap each gene was counted using featureCounts from the subread v2.0.0 package (Liao et al., 2014).

To generate the gene annotations used in featureCounts, we used the previously generated “byexon-gene” and “bysnpgene” annotations (Song et al., 2021) with the following filters: We first filtered out exons or SNPs where more than 10% of reads from diploid and autotetraploid lines map to the incorrect species in NPCs. After removing those exons and SNPs, we then filtered out genes where more than 10% of reads from diploid and autotetraploid lines map to the incorrect species in NPCs when quantifying gene counts from exons separately and from SNPs separately. Finally, we combined exons and SNPs into one annotation where only SNPs that did not overlap exons were included. Genes where more than 10% of reads from diploid and autotetraploid lines map to the incorrect species in NPCs when quantifying gene counts with both exons and SNPs were removed. This resulted in a final annotation containing 484,032 exons and 74,509 SNPs from 99,164 genes.

After sequencing reads were aligned to the composite human-chimpanzee genome, differential gene expression (DE) analysis and allele-specific gene expression (ASE) analysis were performed with DESeq2 using default parameters (Love et al., 2014). We called genes as significant if they had an adjusted *p* < 0.05 after Benjamini-Hochberg FDR correction. Genes were classified by regulatory type based on the following criteria:

- *cis*: significant DE, significant ASE, log_2_(FC) in the same direction between DE and ASE
- *trans*: significant DE, not significant ASE
- *compensatory*: not significant DE, significant ASE

Read counts were also adjusted for feature length to account for differences between feature length in the human and chimpanzee genomes. Results were nearly identical with and without length correction, and only genes that were found as DE or ASE both with and without length correction were considered significant. All visualizations were performed with read counts without length correction. Unless otherwise indicated, a *FC* > 1.5 cut-off was used for all downstream gene set analyses.

### Differential expression analysis of CRISPRi in diploid cells

UMIs were extracted using umi tools v1.1.2 (Smith et al., 2017) and adapter sequences were trimmed using cutadapt v2.5 (Martin, 2011). Sequencing quality was assessed with fastqc v0.11.5 (Andrews et al., 2010). Reads that map to rDNA (https://www.ncbi.nlm.nih.gov/nuccore/U13369.1) were removed, and remaining reads were aligned using STAR v2.7.9a with two-pass mapping (Dobin et al., 2013) to either the hg38 or panTro6 genome, as appropriate. Reads were then de-duplicated based on UMIs using umi tools, and the number of reads that overlap each gene was counted using featureCounts from the subread v2.0.0 package with default parameters (Liao et al., 2014). Differential gene expression analysis was performed with DESeq2 using default parameters (Love et al., 2014). Genes with an adjusted *p* < 0.05 after Benjamini-Hochberg FDR correction were considered to be significant.

### Additional analyses of RNA-seq data

Principal component analysis (PCA) was performed with DESeq2 (Love et al., 2014). Significant gene-set enrichments and gene ontology enrichments (adjusted *p* < 0.05 after Benjamini-Hochberg FDR correction) were determined using the R package clusterProfiler’s gseGO and enrichGO functions respectively (Yu et al., 2012) for the annotation data sets “Biological Process”, “Molecular Function”, and “Cellular Component”. GO enrichments were performed without a fold-change cut-off. The set of analyzed genes was used as the background reference list for gene ontology enrichments. To assess the correlation between two datasets (e.g. Fig. S4E), we used orthogonal distance regression.

*Differential accessibility (DA) and allele-specific accessibility (ASA) analyses of diploids, autotetraploids, and allotetraploids* ATAC-seq data from the human and chimpanzee diploids, autotetraploids, and allotetraploids was aligned to a composite human-chimpanzee genome (hg38 and panTro6) using bwa mem v0.7.17 (Li and Durbin, 2009). Reads with poor alignment or quality were removed (samtools view -F 3844 -q 10) and de-duplicated with samtools v1.3.1 (Li et al., 2009). Bam files were converted to bed files using bedtools v2.26.0 (Quinlan and Hall, 2010), and peaks were called with macs2 v2.1.1.20160309 with the parameters –nomodel –extsize 200 (Zhang et al., 2008). We used reads from diploids and autotetraploids that mapped to the incorrect species as the background for peak calling.

Aligned reads and peaks were then filtered by mapping reads and peaks called in panTro6 to hg38 and vice versa using pslMap from UCSC v363 (Zhu et al., 2007). To generate the chain file used in pslMap, we downloaded panTro6.hg38.all.chain.gz from the UCSC Genome Browser (Navarro Gonzalez et al., 2021) and filtered for orthologous chains as previously described (Turakhia et al., 2020). Reads and peaks were then filtered for those that map uniquely and where the orthologous regions have <5-fold change and <5kb difference in size. Peaks in hg38 coordinates that overlapped ENCODE blacklist regions (Amemiya et al., 2019) or alternate chromosomes were also filtered out. This resulted in reads and peaks in hg38 and panTro6 coordinates for each sample. For allotetraploids, reads and peaks that initially mapped to hg38 were represented separately from those that initially mapped to panTro6.

Reads from diploids and autotetraploids that mapped to the incorrect species were similarly filtered to generate a background mis-mapping file for chimpanzee samples in both hg38 and panTro6 coordinates and for human samples in both hg38 and panTro6 coordinates. We called peaks using macs2 for the background mis-mapping files in hg38 coordinates to identify genomic regions where there were high levels of mis-mapping. Differential accessibility (DA) analysis and allele-specific accessibility (ASA) analysis were performed with DiffBind v3.0.15 (Ross-Innes et al., 2012) for reads and peaks in hg38 coordinates and separately for those in panTro6 coordinates, with the appropriate background mis-mapping files. We analyzed 107,562 ATAC peaks found in at least three samples that could uniquely map to the opposite species. 76,397 of these ATAC peaks did not overlap coding regions. We called peaks as significant if they had an adjusted *p* < 0.05 after Benjamini-Hochberg FDR correction.

Only peaks that were significant with the same direction-of-effect when assessed in both hg38 and panTro6 coordinates were retained. Peaks were classified by regulatory type based on the following criteria:

- *cis*: significant DA, significant ASA, log_2_(FC) in the same direction between DA and ASA
- *trans*: significant DA, not significant ASA
- *compensatory*: not significant DA, significant ASA

For any genome-wide analyses and visualizations, we excluded peaks that overlapped peaks called from background mis-mapping files. In addition, we excluded ATAC peaks overlapping chr21:43377000-43584000 where there is an assembly error in hg38. Principal component analysis (PCA) was performed with prcomp from stats v4.1.3 in R. GREAT was used for gene ontology enrichments (adjusted *p* < 0.05 after Benjamini-Hochberg FDR correction) for the annotation data sets “Biological Process”, “Molecular Function”, and “Cellular Component” (McLean et al., 2010). The set of peaks analyzed by DiffBind were used as the background peak set for GREAT. In Fig. S1E, GREAT enrichments are shown when using all peaks. Similar enrichments are seen when examining only non-coding peaks.

### TF motif analysis of ATAC-seq data

*Cis*- and *trans*-regulated open chromatin regions were separated into those that were human-biased and chimpanzee-biased. For human-biased open chromatin regions, we extracted sequences in hg38 coordinates. For chimpanzee-biased open chromatin regions, we extracted sequences in panTro6 coordinates. Matched background sequences were generated using biasaway v3.3.0 (Khan et al., 2021). PWMSCAN (Ambrosini et al., 2018) was used to scan sequences for motifs from the JASPAR motif database (Castro-Mondragon et al., 2022) to identify potential transcription factor binding sites. To control for false positives, a p-value cutoff of 10^−4^ was used. The presence of motifs was then aggregated and the enrichment of specific motifs compared to matched background sequences and between human-biased and chimpanzee-biased sequences were determined. P-values for enrichment were generated using the proportion test (prop.test in R) and were adjusted for multiple hypothesis testing at 5% FDR with the Benjamini-Hochberg correction. In Fig. 1F, Fig. 4B, Fig. 6B, and Fig. 6C, the adjusted p-value from the comparison between human-biased and chimpanzee-biased sequences was plotted. *Cis*-regulated TFs were highlighted if their motif was significant in the background comparison for the species where the TF is more highly expressed, their motif was significant between human-biased and chimpanzee-biased sequences, and the direction of gene expression change matches the direction of enrichment between human-biased and chimpanzee-biased sequences. *De novo* motif discovery was also performed for human-biased and chimpanzee-biased open chromatin regions with findMotifsGenome.pl from HOMER v4.10 (Heinz et al., 2010).

### Combined analyses of RNA and ATAC sequencing

To compare the proportions of nonsignificant, *cis*-regulated, and *trans*-regulated open chromatin regions, we used the chisquare test at 5% FDR. Open chromatin regions were assigned to genes using GREAT default parameters for all protein-coding genes in the genome or for genes expressed in NPCs. Results were highly concordant between the two sets, and results using only expressed genes are shown in Fig. S3A.

To ask whether there are more human-biased, *cis*-regulated open chromatin regions near human-biased, *cis*-regulated genes when compared to all other genes (and for all other combinations of *cis* and *trans* open chromatin regions and gene), we performed the Kolmogorov–Smirnov test and assessed significance at 5% FDR. Results were similar using all protein-coding genes or all expressed genes, and results from using expressed genes are shown in Fig. S3B.

For the piN RNA-seq and ATAC-seq, we used all RNA-seq and ATAC-seq samples from (Carter et al., 2025) for analyses to maximize power. Although some samples were exposed to KCl stimulation, most species differences in gene expression and chromatin accessibility were independent of the treatment condition (Carter et al., 2025).

### Identifying cis-regulated open chromatin regions for CRISPR inhibition screen

*Cis*-regulated open chromatin regions were linked to *cis*-regulated genes that were either (1) within 500kb or (2) within 1Mb and in the same topologically associated domain (TAD) in human NPCs (Schmitt et al., 2016), human fetal germinal zone (Won et al., 2016), mouse NPCs (Bonev et al., 2017), or NPCs isolated from mouse neocortex (Bonev et al., 2017). Only *cis*-regulated open chromatin regions associated with 1+ *cis*-regulated genes were considered. *Cis*-regulated open chromatin regions were also required to meet the following technical criteria:

- >2kb from any transcription start site (Gencode v40 annotation (Frankish et al., 2019))
- >1kb from any exon (Gencode v40 annotation (Frankish et al., 2019))
- 1+ nearby *cis*-regulated genes are expressed in >20% of cells by scRNA-seq when each cell is sequenced to 40,000 reads - This is the minimal expression level needed to be powered to detect a 25% change in target gene expression.

We then further filtered for *cis*-regulated open chromatin regions that met the following biological criteria:

- Contains 1+ variants that are conserved in marmoset, macaque, gorilla, and chimpanzee and different in humans, suggesting the variant is derived in humans
- Near 1+ *cis*-regulated genes where the expression change between humans and chimpanzees is significantly larger at 10% FDR than the expression change within extant humans as previously described (Starr et al., 2023), suggesting that the human-chimpanzee difference in expression may be functional

Finally, we prioritized *cis*-regulated open chromatin regions that met most or all of the following criteria:

- Near 1+ *cis*-regulated genes with fold change > 1.5
- Near 1+ *cis*-regulated genes where the direction of the gene expression change is consistent between human and chimpanzee NPCs and between human and macaque fetal brain (Zhu et al., 2018), suggesting that the gene expression change is likely derived in humans
- Near 1+ *cis*-regulated genes have pLI = 1 (Lek et al., 2016), suggesting that modest changes in expression may have phenotypic effects
- Near 1+ *cis*-regulated genes implicated in neurodevelopmental disorders (Firth et al., 2009; Leblond et al., 2021)

To design gRNAs, we input the 200bp with the greatest read pile-up from DiffBind in hg38 coordinates to GuideScan (Perez et al., 2017) and filtered for gRNAs with a specificity score > 0.2. If there were not at least three usable gRNAs, we padded the 200bp input up to the size of the ATAC peak in 100-200bp increments to enrich for gRNAs that targeted regions of functionality. We also filtered out peaks that overlapped BstXI or BlpI sites for cloning. We then checked that the exact gRNA sequence is also found in the panTro6 genome at the orthologous location. Only peaks with 3+ gRNAs after filtering were considered.

Peaks that overlapped background mis-mapping regions were manually inspected. Manual inspection revealed that many peaks overlapped background mis-mapping regions only at peak boundaries, and for those regions, read pile-ups suggest that mis-mapping likely had a limited effect. All peaks assessed in the CRISPRi screen passed manual inspection.

### CRISPR inhibition screen analysis

Sequencing reads from gene expression libraries were mapped to hg38 or panTro5 using cellranger v7.0.0 (10X Genomics). The 3’ UTR of *TNIK* is truncated in the panTro5 annotation and was corrected using the orthologous hg38 annotation. gRNAs expressed in each cell were also assigned with cellranger. Downstream analysis was performed with Seurat v4.3.0 (Stuart et al., 2019). Data from NCRM5 was filtered for feature number between 2500-7500, percent mitochondrial reads <10%, and percent ribosomal reads <30%. Data from C3-C was filtered for feature number between 1750-6000, percent mitochondrial reads <10%, and percent ribosomal reads <30%. Data was normalized with SCTranform using 3000 features and regressing out the percent of mitochondrial reads.

We examined the expression of *cis*-regulated genes linked to the tested *cis*-regulated open chromatin regions by comparing gene expression in cells expressing a test gRNA compared to cells expressing non-targeting control (NTC) gRNAs. Comparisons were performed for test gRNA and potential target genes where there were >10 cells expressing the test gRNA, >1 cell expressing the test gRNA and the gene of interest, >1 cell expressing NTC gRNAs and the gene of interest, and where 5% of cells expressing either the test gRNA or NTC gRNAs expressed the gene of interest. We performed a likelihood ratio test on a generalized linear model where the counts were modeled using a negative binomial distribution as previously described (Gasperini et al., 2019). The generalized linear model was fit with the percent of mitochondrial RNA and percent of ribosomal RNA as additional variables.

We then performed the same differential expression analysis for each NTC gRNA and each tested gene, and calculated empirical p-values based on this distribution as previously described (Gasperini et al., 2019). Nonrandom assortment of gRNAs in cells was assessed with Fisher’s exact test, and statistical tests between NTC gRNAs and genes near a nonrandomly assorted test gRNA were excluded. Empirical p-values were then Benjamini-Hochberg corrected, and significance was assessed at 5% FDR. We also performed this analysis for all genes within 1Mb of the gRNA target sites. Further, we compared results when examining all cells or only cells that expressed a single gRNA. Although we were underpowered to assess significance in most cases when only considering cells that expressed a single gRNA, the effect sizes were highly concordant.

### CUT&Tag sequencing analysis

CUT&Tag sequencing data was processed as described above for the ATAC-seq data with the following differences. Prior to mapping, adapters were trimmed with cutadapt v2.5 (Martin, 2011) and sequencing quality was determined with fastqc v0.11.5 (Andrews et al., 2010). Peak calling with macs2 was performed with matched IgG samples as the background set and the following parameters: –nomodel –shift -100 – extsize 200. Peak calling was also performed for all mapped reads for each TF of interest using macs2 to get a list of peaks as the input regions for DiffBind. Only non-coding peaks are visualized and described in Fig. 4, Fig. 5, and associated text.

*De novo* motif discovery was performed on called peaks using hg38 coordinates and findMotifsGenome.pl from HOMER v4.10 (Heinz et al., 2010). The enrichment of CUT&Tag reads at predicted binding sites from the ATAC-seq data (from PWM-SCAN as described above) was determined using makeTagDirectory and annotatePeaks.pl from HOMER. GREAT was used for gene ontology enrichments (adjusted *p* < 0.05 after Benjamini-Hochberg FDR correction) for the annotation data sets “Biological Process”, “Molecular Function”, and “Cellular Component” (McLean et al., 2010). The whole genome was used as the background for GREAT. There were no GREAT enrichments for *trans*-regulated MAZ binding sites.

### Analysis of luciferase assays

Nanoluciferase activity was normalized to firefly luciferase activity for each well. Luciferase activity measurements were performed with 3 replicates per experiment and the mean value was used as the luciferase activity measurement for that experiment. Luciferase activity measurements were further normalized to the luciferase activity of the null condition (pNL3.2 or 0µM KY-05009). For the Wnt reporter assay, we assessed whether 50µM KY-05009 significantly decreased Wnt reporter activity using the one-sided wilcoxon rank-sum test at 5% FDR. For enhancer reporter assays, each test vector was first assessed for whether it acted as an enhancer compared to the empty vector (pNL3.2) using the one-sided wilcoxon rank-sum test. For test regions where the human or chimpanzee sequence acted as an enhancer at 5% FDR, luciferase activity was compared between the human and chimpanzee sequences using the paired wilcoxon rank-sum test. Test regions where the human and chimpanzee sequences have an adjusted *p* < 0.05 after Benjamini-Hochberg FDR correction were considered to be significant.

### Analysis of qPCR

To determine if different CRISPRi gRNAs had different effects on *TNIK* expression (Fig. 2F, Fig. S2E), *TNIK* expression was first normalized to *GAPDH* expression within each sample. Normalized *TNIK* expression was then further normalized to the mean *TNIK* expression when NPCs were infected with NTC gRNAs in each experiment and in each cell line. An analysis of variance (aov) model was fit for normalized *TNIK* expression to cell line, experiment, and gRNA category (NTC, targeting RE-*TNIK*, targeting RE2-*TNIK*, targeting the *TNIK* promoter, or targeting both RE-*TNIK* and RE2-*TNIK*), and statistical significance was assessed with simultaneous tests for generalized linear hypotheses using multiple comparisons of means and Tukey contrasts.

### Intersection with HARs and HAQERs

HAR sets are from (Girskis et al., 2021; Keough et al., 2023; Bi et al., 2023). HAQERs are from (Mangan et al., 2022). HAR and HAQER sets were intersected with all, *cis, trans*, and *compensatory* open chromatin regions using bedtools v2.30.0 (Quinlan and Hall, 2010). The proportion of all, *cis, trans*, or *compensatory* open chromatin regions that overlap with HARs or HAQERs was calculated and divided by the proportion of all open chromatin regions that overlap with HARs or HAQERs. The chi-square test was used to assess statistical significance at 5% FDR.

### Analysis of MEA recordings

The AXIS Neural Metric Tool was used to generate CSV files containing recording metadata and neuronal activity metrics. These metrics were analyzed in R as described in (Carter et al., 2025). For analysis, three NTC gRNAs were combined, and three gRNAs targeting the *POU3F2* promoter were combined.

