## Supplemental Figures for "Human-chimpanzee tetraploid system defines mechanisms of species-specific neural gene regulation"

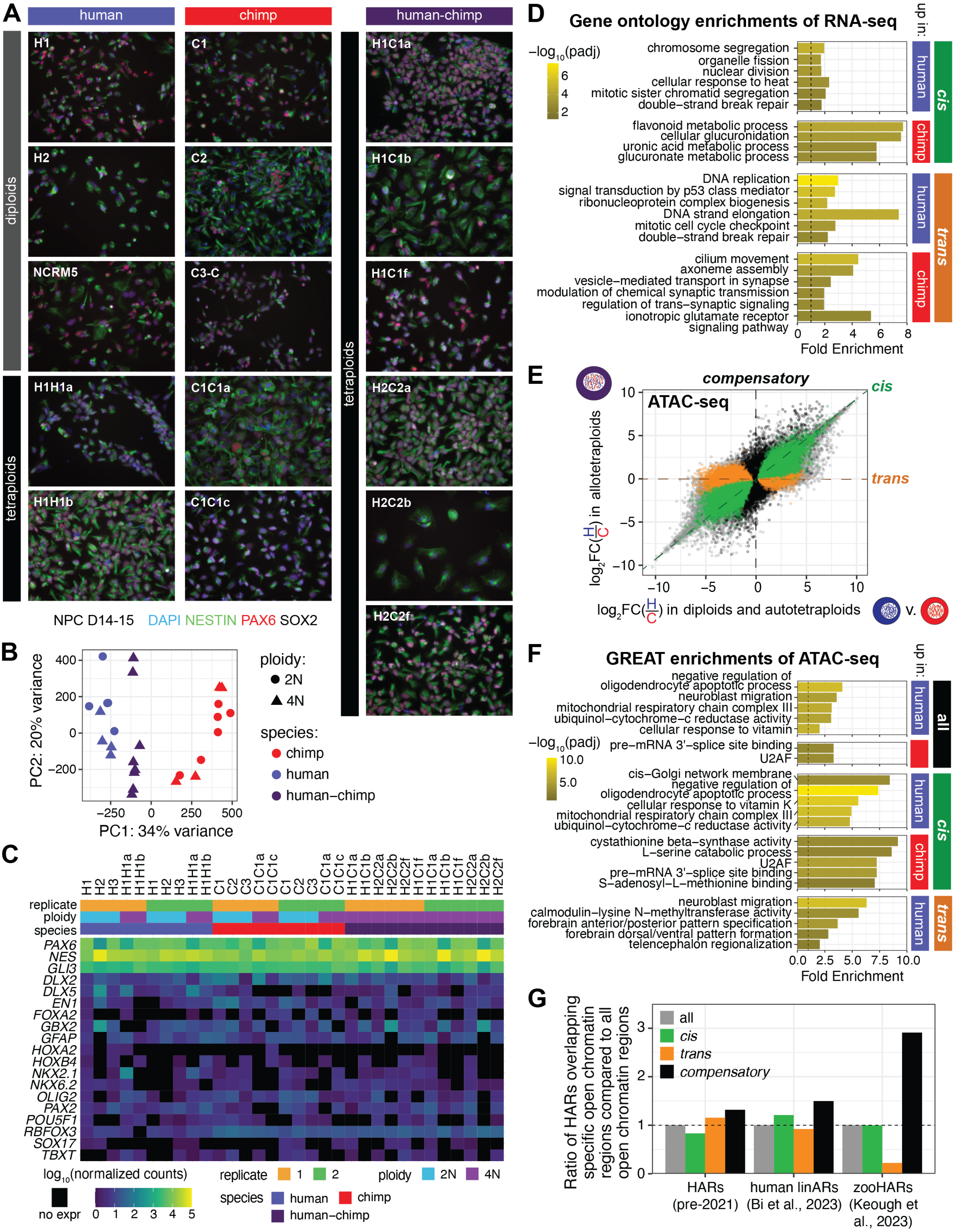


**Figure S1: Marker gene expression and gene ontology enrichments for RNA-seq and ATAC-seq in NPCs, Related to Fig. 1.** (Continued on next page)

(Continued from previous page) (A) NESTIN, PAX6, and SOX2 immunofluorescence staining of NPCs differentiated for 14-15 days from diploid, autotetraploid, and allotetraploid iPSC lines. H2C2a is also shown in Fig. 1B. (B) PCA of ATAC-seq shows separation by species. (C) Heatmap of normalized gene expression counts for marker genes from RNA-seq data. (D) Representative gene ontology enrichments of RNA-seq data. (E) ATAC-seq identifies *cis*-regulated (green), *trans*-regulated (orange), and *compensatory* (black) open chromatin regions. (F) Representative GREAT enrichments of ATAC-seq data. There were no terms enriched for chimpanzee-biased, *trans*-regulated open chromatin regions. (G) Ratio of the proportion of HARs identified prior to 2021 (Girskis et al., 2021), human linARs (Bi et al., 2023), and zooHARs (Keough et al., 2023) overlapping all, *cis*-regulated, *trans*-regulated, or *compensatory* open chromatin regions compared to the proportion of HAR subsets overlapping all open chromatin regions.

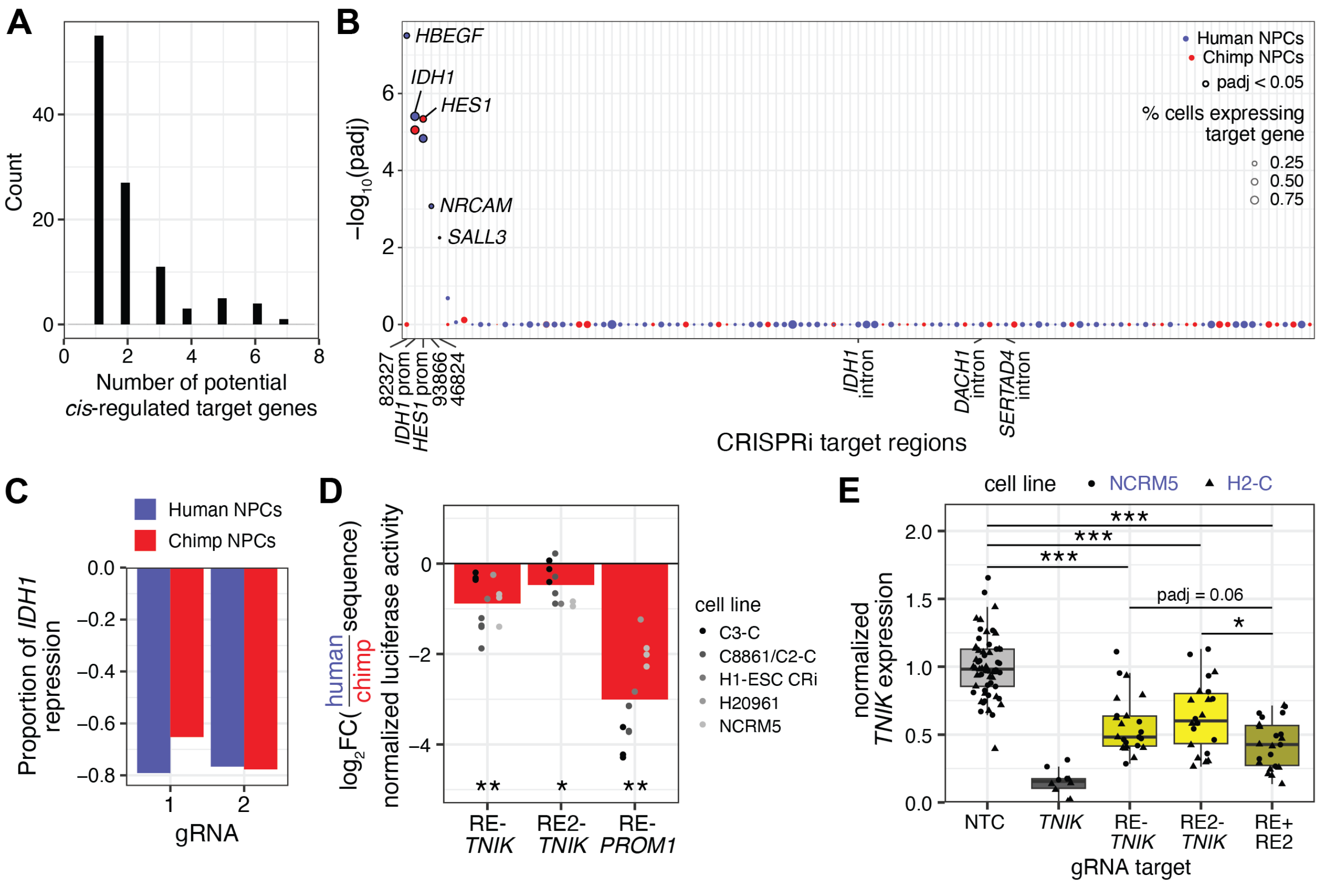


**Figure S2: CRISPR inhibition screen in NPCs identifies *cis*-regulated enhancers of nearby genes, Related to Fig. 2.** (A) Histogram of number of potential *cis*-regulated target genes near each *cis*-regulated open chromatin region assessed in the CRISPRi screen. (B) Adjusted p-values for the effect of targeting each *cis*-regulated open chromatin region on a nearby gene that is not *cis*-regulated in human NPCs (blue) or chimpanzee NPCs (red). Only the nearby gene with the highest p-value is shown. Significant hits are outlined in black and labeled with their target gene. Positive controls targeting promoters (*IDH1* prom and *HES1* prom) and negative controls targeting introns are also labeled. (C) Proportion of *IDH1* repression for the two gRNAs targeting the *IDH1* promoter in human and chimpanzee NPCs. (D) Fold change between luciferase activities of human and chimpanzee sequences for RE-*TNIK*, RE2-*TNIK*, and RE-*PROM1*. (E) qPCR for *TNIK* expression in NPCs differentiated from 2 human lines (NCRM5 and H2-C) and infected with a combination of a NTC gRNA and gRNAs targeting the *TNIK* promoter, RE-*TNIK*, or RE2-*TNIK* (Materials and Methods). Note that statistical analysis was performed with data from both human lines and chimpanzee lines (Fig. 2F). *: $padj < 0.05$; **: $padj < 0.01$; ***: $padj < 0.001$.

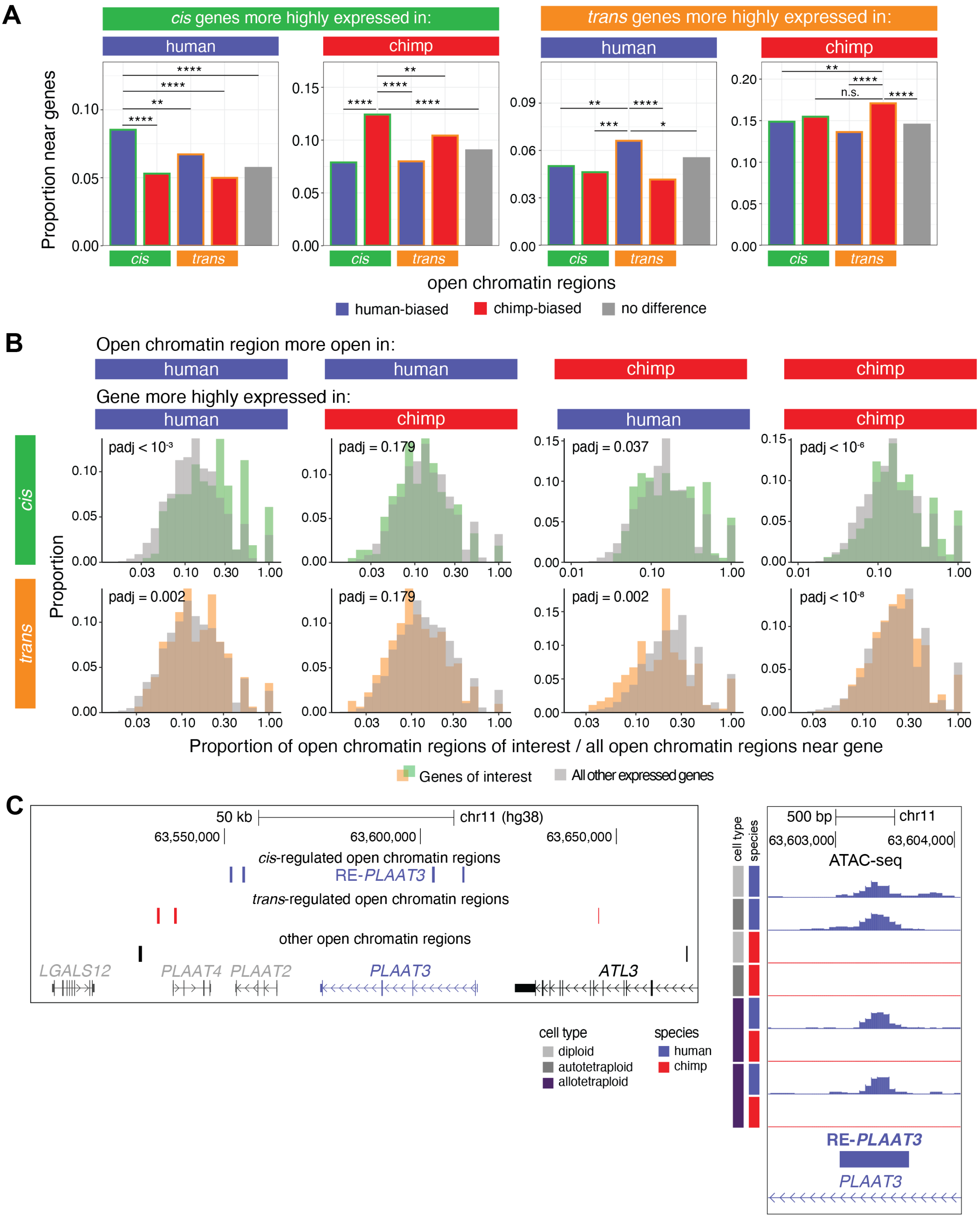


**Figure S3: Enrichment of *cis*- and *trans*-regulated open chromatin regions near *cis*- and *trans*-regulated genes, Related to Fig. 2.** (A) Proportion of open chromatin regions with different types of regulation near genes with different types of regulation. n.s.: $padj > 0.05$; *: $padj < 0.05$; **: $padj < 0.01$; ***: $padj < 0.001$; ****: $padj < 0.0001$. (Continued on next page)

(Continued from previous page) (B) Proportion of *cis*- and *trans*-regulated open chromatin regions near each gene compared to all open chromatin regions near that gene for *cis*-regulated genes, *trans*-regulated genes, and all expressed genes. Genes without any nearby open chromatin regions of interest ($x - value = 0$) are not plotted, but were included in the statistical analyses (Materials and Methods). For categories with the same direction of effect between open chromatin regions and genes (columns 1 and 4), there were on average 9.3% less genes without any nearby open chromatin regions of interest ($x - value = 0$) for the genes of interest compared to all other expressed genes. For categories with the opposite direction of effect (columns 2 and 4), there were on average 2.6% more genes without any nearby open chromatin regions of interest ($x - value = 0$) for the genes of interest compared to all other expressed genes. For instance, for *trans* - open chromatin region more open in chimpanzee - gene more highly expressed in chimpanzee, 27.1% of genes of interest had no nearby open chromatin regions of interest compared to 38.6% of other expressed genes. (C) *PLAAT3* is near nine open chromatin regions (left). All *cis*-regulated open chromatin regions in this interval are human-biased, and all *trans*-regulated open chromatin regions in this interval are chimpanzee-biased. *LGALS12*, *PLAAT4*, and *PLAAT2* are not expressed in NPCs. *ATL3* is expressed but is not species-differential. Read pile-ups from ATAC-seq are shown for representative diploid, autotetraploid, and allo-tetraploid NPCs for the human-biased, *cis*-regulated open chromatin region RE-*PLAAT3* (right).

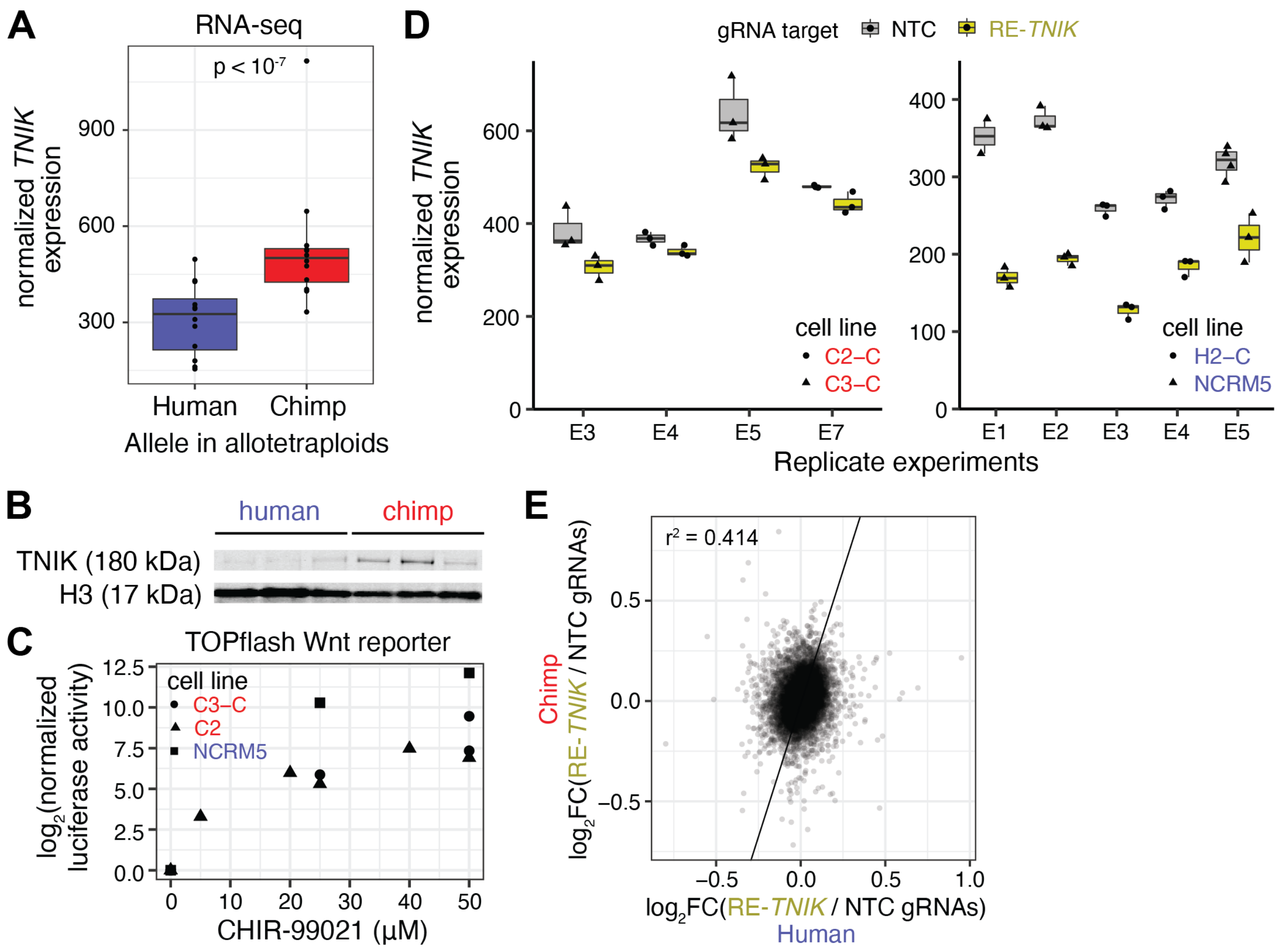


**Figure S4: RE-*TNIK* is a species-differential enhancer that affects cell cycle gene expression in NPCs, Related to Fig. 3.** (A) Allele-specific expression of *TNIK* in allotetraploid NPCs (Table S1). (B) Western blot of NPCs differentiated from three human diploid lines (H1, H2, NCRM5) and three chimpanzee diploid lines (C1, C2, C3) for TNIK and H3. (C) Increasing concentrations of CHIR-99021 increased luciferase activity from the Wnt reporter TOPflash in NPCs differentiated from two chimpanzee cell lines (C2, C3-C) and one human cell line (NCRM5). (D) Normalized gene counts of *TNIK* in NPCs targeted with either NTC gRNAs (gray) or gRNAs targeting RE-*TNIK* (yellow). NPCs were differentiated from two chimpanzee lines (C2-C, C3-C) and two human lines (H2-C, NCRM5). (E) Scatterplot of fold changes for expressed genes between NPCs gRNAs targeting RE-*TNIK* and NPCs infected with NTC gRNAs in human NPCs compared to chimpanzee NPCs.

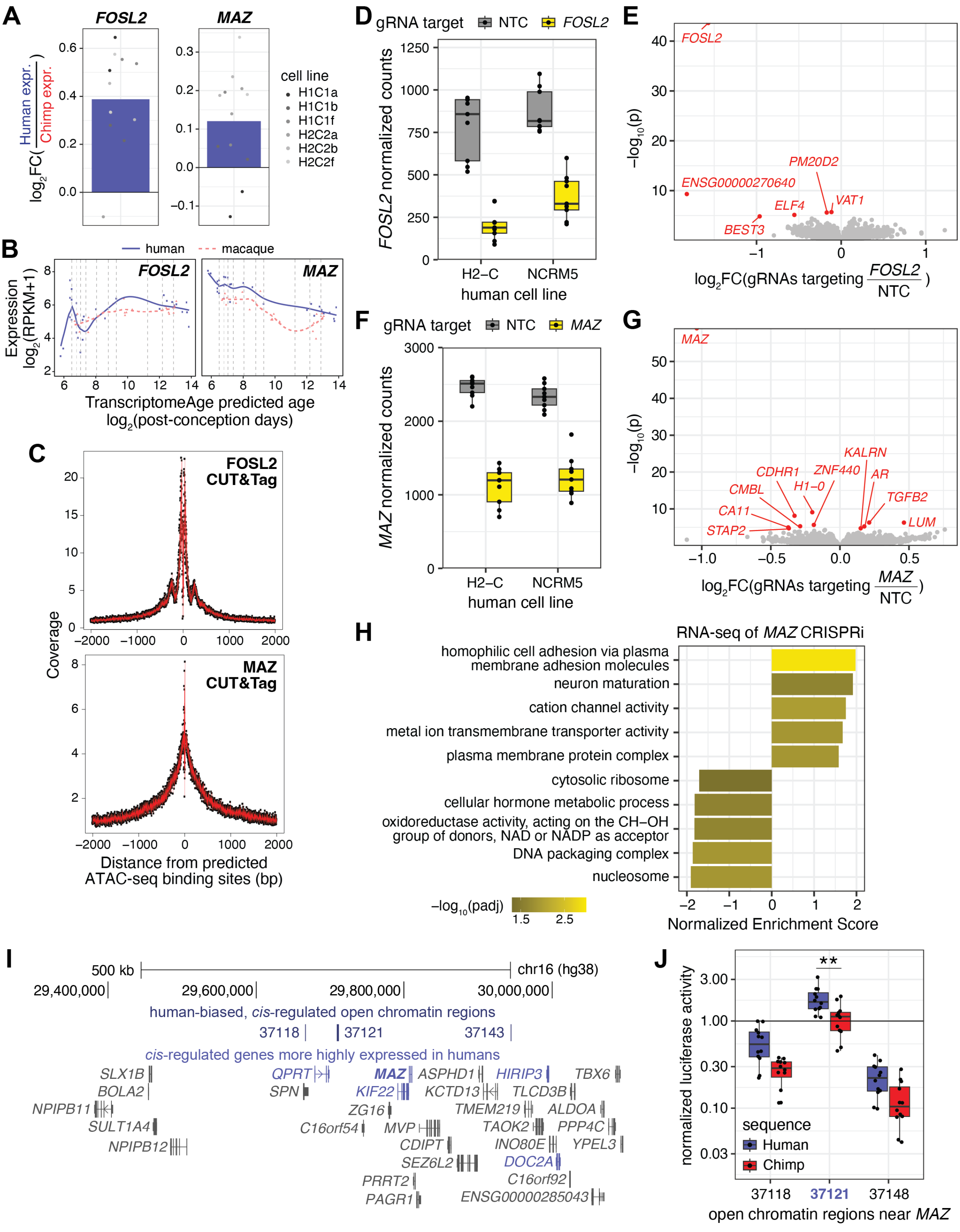


**Figure S5: Characterization of *cis*-regulated TFs, FOSL2 and MAZ, in NPCs, Related to Fig. 4 and Fig. 5.** (A) Fold change between *FOSL2* or *MAZ* expression from human and chimpanzee alleles in allotetraploid NPCs. (Continued on next page)

(Continued from previous page) (B) Expression of *FOSL2* and *MAZ* during human and macaque brain development in the dorsolateral prefrontal cortex from (Zhu et al., 2018). Gene expression profiles are similar across other brain regions assessed in (Zhu et al., 2018). (C) FOSL2 and MAZ binding is enriched at binding sites predicted from ATAC-seq data. (D) Normalized gene counts of *FOSL2* in human NPCs targeted with either NTC gRNAs (gray) or gRNAs targeting the *FOSL2* promoter (yellow). (E) Volcano plot of differential gene expression between human NPCs infected with gRNAs targeting the *FOSL2* promoter or NTC gRNAs. Genes significant at 5% FDR are in red. (F) Normalized gene counts of *MAZ* in human NPCs targeted with either NTC gRNAs (gray) or gRNAs targeting the *MAZ* promoter (yellow). (G) Volcano plot of differential gene expression between human NPCs infected with gRNAs targeting the *MAZ* promoter or NTC gRNAs. Genes significant at 5% FDR are in red. (H) Gene-set enrichment analysis of RNA-seq of human NPCs infected with gRNAs targeting the *MAZ* promoter compared to those infected with NTC gRNAs. (I) TAD containing *MAZ* contains three human-biased, *cis*-regulated open chromatin regions, as well as four other *cis*-regulated genes with higher expression in humans. (J) Luciferase enhancer activity of the human and chimpanzee sequences of the three *cis*-regulated open chromatin regions near *MAZ*. **: $padj < 0.01$.

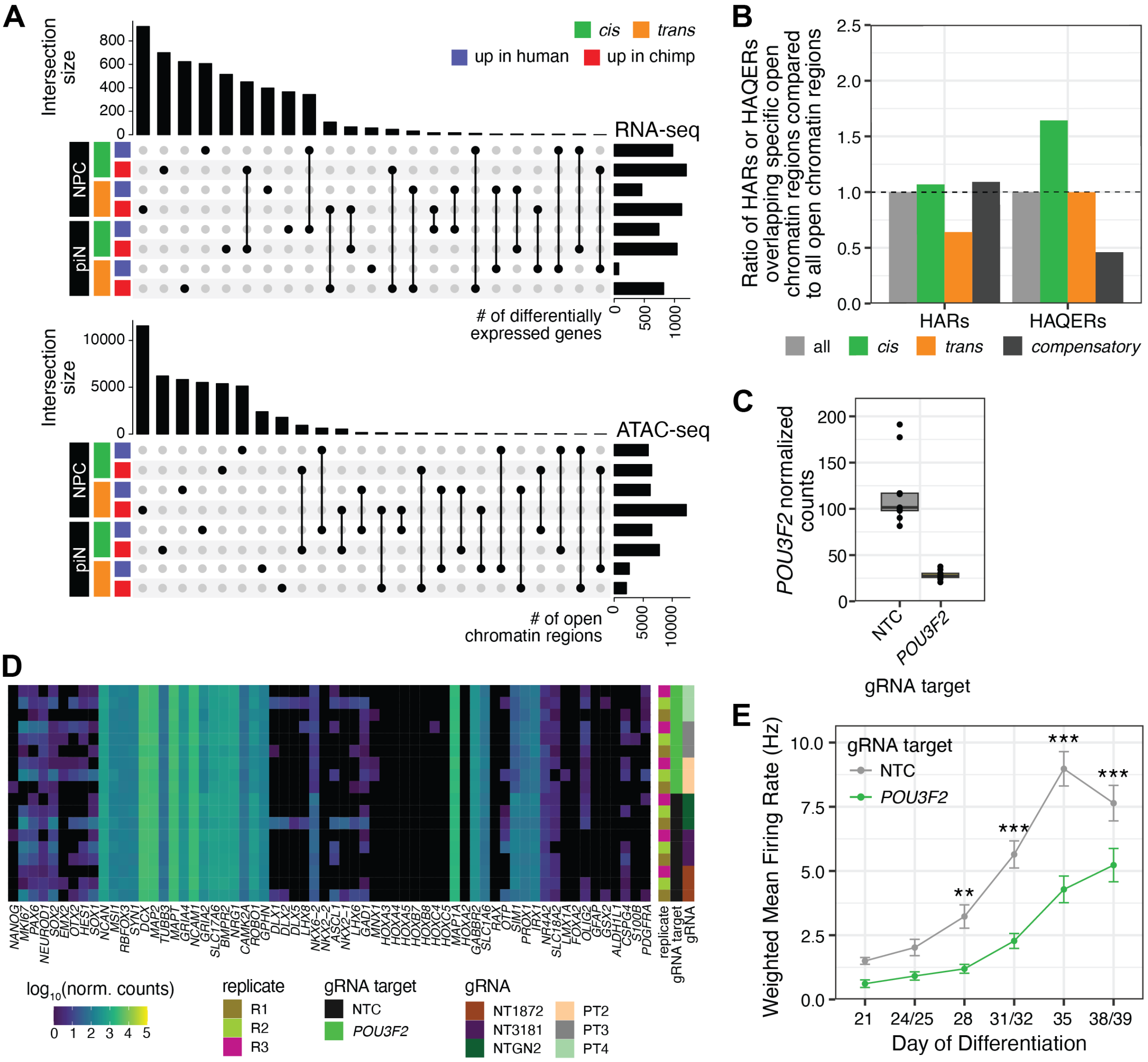


**Figure S6: Gene regulatory differences between human and chimpanzee piNs, Related to Fig. 6.** (A) Upset plots comparing RNA-seq (top) and ATAC-seq (bottom) between NPCs and piNs, separated by regulation type and species bias. (B) Ratio of the proportion of HARs or HAQERs overlapping all, *cis*-regulated, *trans*-regulated, or *compensatory* open chromatin regions compared to the proportion of HARs or HAQERs overlapping all open chromatin regions. *Cis* to all open chromatin regions for HAQERs: $p = 0.028$. (C) Normalized gene counts of *POU3F2* in piNs targeted with either NTC gRNAs (gray) or gRNAs targeting the *POU3F2* promoter (green) for piNs differentiated from the human line NCRM5. (D) Normalized gene counts of marker genes from (Nehme et al., 2018) in piNs targeted with either NTC gRNAs or gRNAs targeting the *POU3F2* promoter. (E) Weighted mean firing rate of MEA recordings of piNs differentiated from the human line H2-C that were infected with gRNAs targeting the *POU3F2* promoter (green) or with NTC gRNAs (gray). Mean ± SEM is plotted. **: $padj < 0.01$; ***: $padj < 0.001$.

**Supplemental Tables**

Table S1: **RNA-seq analysis of human and chimpanzee diploid, autotetraploid, and allotetraploid NPCs**
Table S2: **ATAC-seq analysis of human and chimpanzee diploid, autotetraploid, and allotetraploid NPCs**
Table S3: **CRISPRi screen of *cis*-regulated open chromatin regions in NPCs**
Table S4: **RNA-seq analysis of CRISPRi targeting RE-*TNIK***
Table S5: **TF enrichments at *trans*-regulated open chromatin regions, FOSL2 and MAZ CUT&Tag**
Table S6: **RNA-seq analysis of CRISPRi targeting FOSL2 or MAZ**
Table S7: **TF enrichments at species-biased open chromatin regions in piNs**
Table S8: **RNA-seq analysis of CRISPRi targeting POU3F2**
Table S9: **Cell lines and oligos used in this study**
